# A multiplex CRISPR interference tool for virulence gene interrogation in an intracellular pathogen

**DOI:** 10.1101/2020.06.17.157628

**Authors:** Nicole A. Ellis, Byoungkwan Kim, Matthias P. Machner

**Affiliations:** Division of Molecular and Cellular Biology, Eunice Kennedy Shriver National Institute of Child Health and Human Development, National Institutes of Health, Bethesda, Maryland, 20892 United States of America

## Abstract

In the absence of target cleavage, catalytically inactive dCas9 imposes transcriptional gene repression by sterically precluding RNA polymerase activity at a given gene to which it was directed by CRISPR (cr)RNAs. This gene silencing technology, referred to as CRISPR <u>i</u>nterference (CRISPR<u>i</u>), has been employed in various bacterial species to interrogate genes, mostly individually or in pairs. Here, we developed a multiplex CRISPRi platform in the pathogen *Legionella pneumophila* capable of silencing up to ten genes simultaneously. Constraints on precursor-crRNA expression by Rho-dependent transcription termination were overcome by combining a strong processive promoter with a *boxA* element upstream of a repeat/spacer array. Using crRNAs directed against virulence protein-encoding genes, we demonstrated that CRISPRi is fully functional not only during growth in axenic media, but also during macrophage infection, and that gene depletion by CRISPRi fully recapitulated the growth defect of deletion strains. Importantly, by altering the position of crRNA-encoding spacers within the repeat/spacer array, our platform achieved the gradual depletion of targets that was mirrored by the severity in phenotypes. Multiplex CRISPRi thus holds great promise for probing large sets of genes in bulk in order to decipher virulence strategies of *L. pneumophila* and other bacterial pathogens.

## Introduction

Clustered regularly interspaced short palindromic repeats (CRISPR)-Cas gene editing technologies have recently arisen as a mechanism for both fast and targeted gene manipulation in a variety of systems (1-3). CRISPR-Cas-based genetic tools are derived from components of naturally occurring CRISPR systems found in 87% of archaea and 45% of bacteria genomes surveyed (Crisprfinder; http://crispr.i2bc.paris-saclay.fr/Server/). In these organisms, CRISPR regions serve as an adaptive immune system where fragments of foreign DNA elements such as viruses, transposons, or plasmids are incorporated into the chromosome as unique spacers (S) separated by identical repeats (R) to serve as a immunological memory of past infections (1, 4, 5). RNA transcribed from these repeat/spacer regions, known as the precursor-CRISPR (cr)RNA, is processed into short fragments by RNase III to make individual crRNAs (6). Upon re-infection, crRNAs, together with a trans-activating crRNA (tracrRNA), specifically direct, through base-pairing with the target, a protein (or contingent of proteins) with oligonucleotide cleavage capabilities to the foreign DNA or RNA elements for destruction (7-9). Importantly, the spacers themselves are protected from self-targeting by the crRNAs that they encode as they are missing the protospacer-adjacent motif (PAM) (10), a short DNA element found directly downstream of the complementary target sequence on the non-target strand that is required by the surveillance complex to cut.

In the simplest CRISPR-Cas system, Type II, only a single protein known as Cas9 is required for crRNA-guided DNA cleavage, lending it to be the most developed as a genetic tool (11-13). In bacteria, this system has been adapted for gene silencing by making two simple changes: First, the gene encoding Cas9 was replaced with a catalytically inactive variant of Cas9, called deactivated Cas9 (dCas9), in which the two nuclease domains, RuvC-like and HNH, have been mutated, thus preventing DNA cleavage (14, 15); and second, by designing arrays in which the spacer sequence(s), that are typically directed against invading DNA elements, have been replaced with sequences complementary to the bacterium’s own genes. These self-targeting crRNAs form a surveillance complex together with the tracrRNA and with dCas9 which, in the absence of cleavage, imposes transcriptional gene repression by sterically precluding RNA polymerase activity at the gene to which the complex was directed, a technology referred to as CRISPR interference (CRISPRi) (14, 16). Gene silencing is advantageous to the study of essential genes that are otherwise intolerable to deletion, as well as for interrogating genes of interest without laborious null strain construction (17). The recent development of mobile-CRISPRi has allowed for gene silencing to be performed in a number of gamma proteobacteria and Bacillales Firmicutes (18), yet this and earlier studies, which implemented CRISPRi (19), nearly always silenced only one gene or pairs of genes, with few exceptions (20-23). To our knowledge, no group has fully exploited or probed the natural multiplex capability of CRISPR repeat/spacer arrays as a gene silencing tool in bacterial systems.

Here, we established an adaptable multiplex CRISPRi platform capable of silencing up to ten genes simultaneously (10-plex). We demonstrate that this platform can be applied to study bacterial virulence genes not only in axenic media but, importantly, also under disease-relevant conditions such as the infection of macrophages by the intracellular pathogen *Legionella pneumophila*, thus adding CRISPRi to our toolbox for efficiently studying genetically less tractable human pathogens.

## Results

### Equipping *L. pneumophila* with a CRISPRi system

To study the potential of multiplex CRISPRi, we selected the model organism *L. pneumophilia*, the causative agent of Legionnaires’ pneumonia or a milder form of the disease called Pontiac fever (24, 25). The common laboratory strain *L. pneumophila* Philadelphia-1 (Lp02) does not bear an intrinsic CRISPR-Cas system (though some environmental *L. pneumophila* isolates do (26)). Thus, we exogenously introduced the genes encoding the three main components of a CRISPRi platform: a Cas protein, crRNAs, and the tracrRNA (Figure 1A). The *Streptococcus pyogenes* dCas9-encoding sequence was inserted into the chromosomal *thyA* locus (*lpg2868*) as this *L. pneumophila* gene was already disrupted by a mutation in this strain background (27), creating Lp02(*dcas9*). Furthermore, *dcas9* was placed under the control of the tetracycline-inducible *tet* promoter (P_*tet*_) rather than its native *S. pyogenes* promoter, allowing the degree of gene repression to reflect the level of anhydrous tetracycline (aTC) added to the system to induce *dcas9*, as others have done before (14). The crRNA-encoding repeat/spacer arrays were provided on plasmids, as was the target-independent (invariable) tracrRNA-encoding sequence. Construction of these single CRISPRi constructs is described in the Methods section and depicted in Figure S1A. Although single guide (sg)RNAs, chimeras of the tracrRNA and crRNA, are convenient for silencing individual genes as to circumvent the need for processing precursor-crRNA into a mature crRNAs (6), the tracrRNA-encoding sequences would become repetitive within longer CRISPR arrays. Instead, the task of processing crRNA precursors was seemingly executed efficiently by the intrinsic *L. pneumophila* RNAse III (*lpg1869*) in our system (see below).

**Figure 1:**
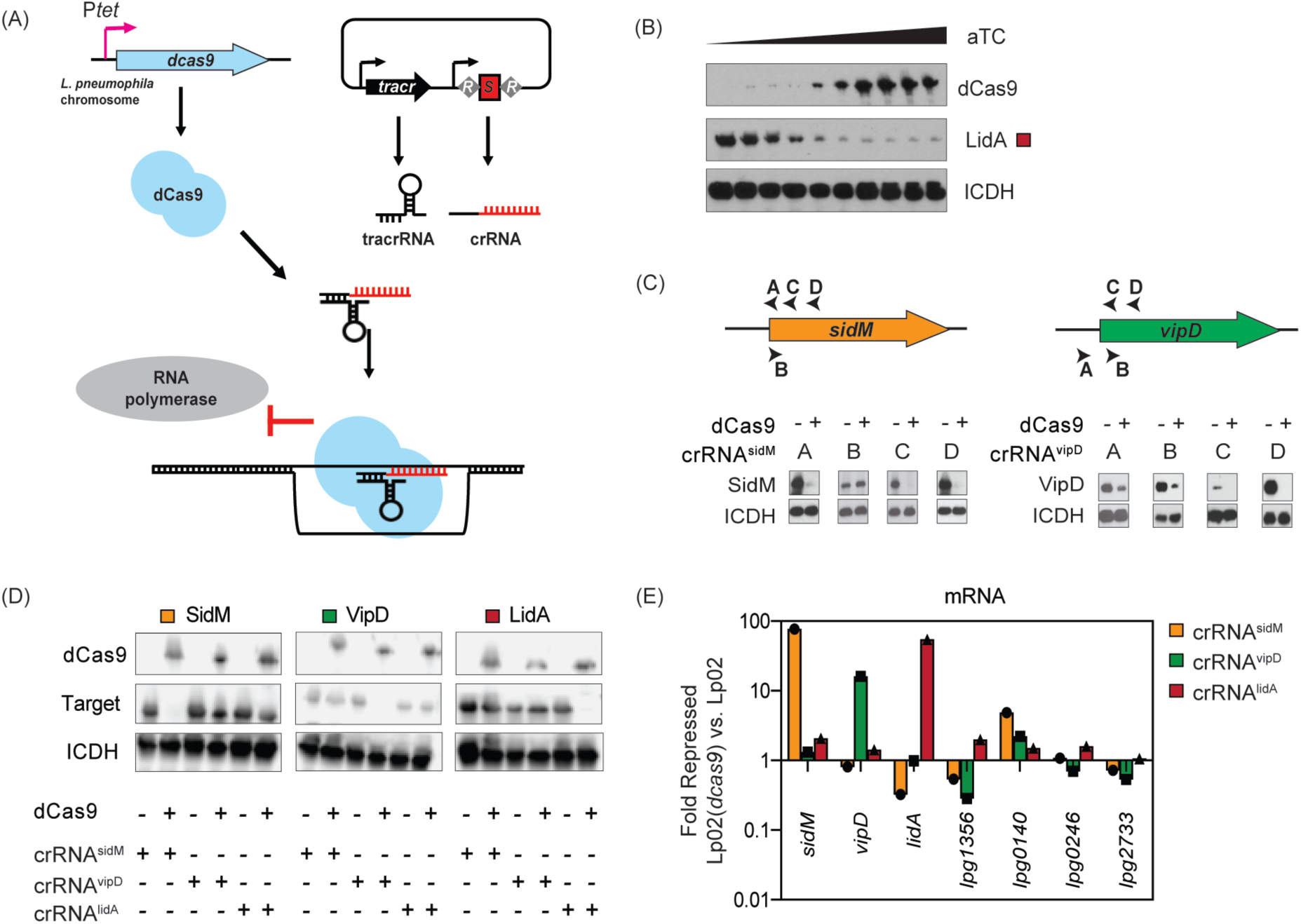
CRISPRi is adaptable to *L. pneumophila*. (A) Schematic of the *L. pneumophila* single-plex CRISPRi platform. Chromosome-encoded dCas9 and plasmid-expressed tracrRNA and crRNA assemble into a CRISPRi complex. The crRNA directs the complex to the target gene through base pairing. Gene expression is repressed through sterically precluding RNA polymerase. R indicates repeats and S indicates spacers of the crRNA-encoding repeat/spacer array. (B) Immunoblot analysis showing the protein level of dCas9 in response to aTC-dependent induction of P_*tet*_-*dcas9*, leading to decreased protein levels of the crRNA^lidA^ target. Isocitrate dehydrogenase (ICDH) serves as a loading control. aTC concentrations are 0, 0.15, 0.3, 0.63, 1.23, 2.5, 5, 10, 20, and 80 ng/mL. (C) Immunoblots reveal the efficiency of target repression at different crRNA target sites (arrow heads A-D) for two genes, *sidM* and *vipD* (+20 ng/mL aTC). ICDH serves as a loading control. (D,E) Strains expressing crRNA^sidM^ (site C), crRNA^vipD^ (site C), and crRNA^lidA^ were surveyed for CRISPRi specificity at both the protein (D) and mRNA (E) level (+40 ng/mL aTC). ICDH serves as an immunoblot loading control. qPCR was used to measure mRNA levels of *sidM, vipD, lidA* and four randomly chosen control genes in Lp02(*dcas9*) and Lp02 strains. Fold repression was determined using ΔΔC_T_. A value of 1 indicates that the gene level was the same in the Lp02(*dcas9*) and Lp02 strains.

Targeting individual genes with single crRNA arrays served as proof-of-concept for introducing a foreign CRISPR-Cas9 platform into our model organism *L. pneumophila*. These initial experiments were performed on *L. pneumophila* strains grown in axenic culture while individually targeting three virulence factor, or “effector”, -encoding genes with crRNA towards *lidA* (*lpg0940*; crRNA^lidA^), *sidM* (*lpg2464*; crRNA^sidM^), and *vipD* (*lpg2831*; crRNA^vipD^), since antibodies directed against their encoded gene products were available. First, we confirmed that a gradual enhancement in the concentration of the inducer aTC lead to increased levels of dCas9 and, consequently, decreased levels of LidA, the product of the target *lidA*, indeed creating a CRISPRi system capable of tunable gene silencing (Figure 1B). Next, we assessed how the location of the crRNA annealing site on the target gene (strand, proximity to start codon) affects gene silencing efficiency. In agreement with other studies (12, 14, 28), we found that targeting a 30 base pair sequence on the coding strand, near the start codon, upstream of a template strand PAM (“NGG”, where “N” is any nucleobase followed by two guanine (“G”) nucleobases, (29)) proved most efficient in decreasing protein levels by immunoblot analysis (Figure 1C, arrowheads C and D).

Lastly, to confirm that crRNAs designed in this manner did not indiscriminately affect gene expression, we monitored protein (Figure 1D) and mRNA (Figure 1E) levels of unrelated genes. Using immunoblot analyses and quantitative polymerase chain reaction (qPCR) we found that, while the intended target genes’ protein and mRNA levels were dramatically reduced by CRISPRi, none of the other surveyed genes chosen at random were repressed by crRNA expression. Together, these results confirmed that the aTC-inducible *dcas9* and the plasmid-expressed crRNA and tracrRNA formed a functional CRISPRi surveillance complex in *L. pneumophila* that was tunable, efficient, and target-specific.

### A multiplex CRISPRi platform allows for the simultaneous repression of several genes

Most processes in bacteria, including pathogenesis, rely on the combined activity of multiple gene products. With over 300 putative effectors, *L. pneumophila* encodes one of the largest known virulence arsenals of any bacterial pathogen (30, 31). We thus explored if our CRISPRi platform could be adapted to simultaneously repress more than one gene at a time. In this new multiplex CRISPRi format, expression of *S. pyogenes dcas9* from the *L. pneumophila* chromosome and the expression of the *S. pyogenes* tracrRNA from the plasmid backbone remained unchanged. Alterations were made to the method of crRNA expression: First, several unique crRNA-encoding spacer sequences, labelled *S1* (for the most proximal spacer) to *S10* (for the most distal), were sequentially placed between identical repeat sequences intrinsic to the *S. pyogenes* CRISPSR-Cas9 system (Figure 2A). In nature, the number of spacers within a CRISPR array can vary, ranging from a few to several dozen or even hundreds (32). Second, expression of the repeats and spacers was placed under the regulation of P_*tet*_, the same aTC-inducible promoter used to express *dcas9* from the *L. pneumophila* chromosome. We had found in experiments on Lp02(*dcas9*) grown in axenic culture that using the native *S. pyogenes* promoter (P_*native*_) to express the multiplex CRISPR (MC) repeat/spacer arrays proved insufficient in repressing more than three to four targets, presumably due to inefficient RNA polymerase processivity driven from a weaker promoter (Figure 2B). Incorporation of P_*tet*_ drove expression up to at least spacer *S9*, as confirmed by immunoblot analysis of the protein levels of LidA whose encoding gene was targeted by crRNAs expressed from the most distal spacer positions (Figure 2B).

**Figure 2:**
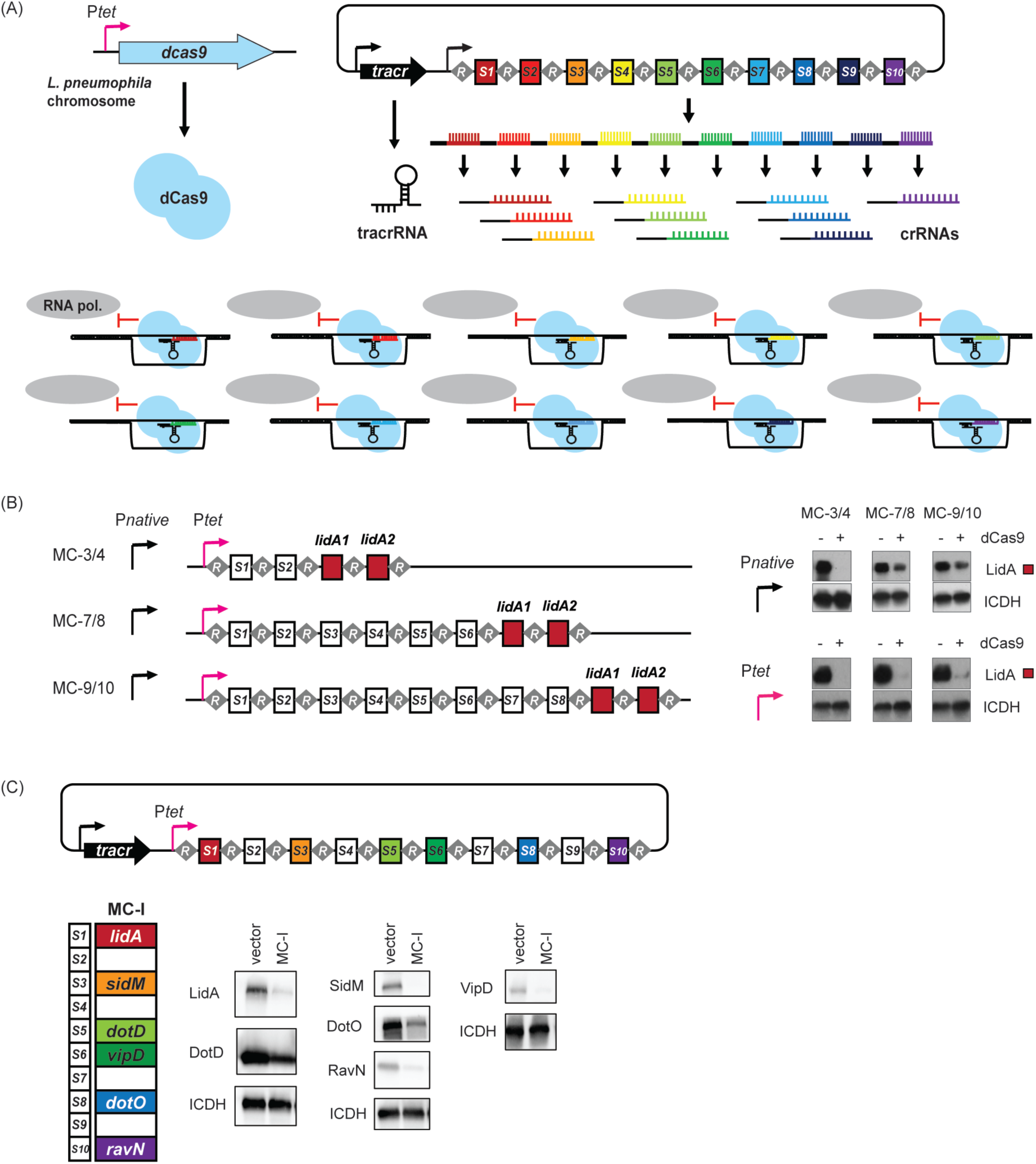
Repeat/spacer arrays facilitate multiplex gene silencing. (A) Schematic representation of the theory of multiplex CRISPRi. A series of repeats, R, and spacers, *S1*-*S10*, are expressed as a single precursor-crRNA. Upon processing, individual crRNAs come together with a tracrRNA and dCas9 to simultaneously target ten unique genes for silencing. (B) <u>M</u>ultiplex <u>C</u>RISPR (MC) repeat/spacer array constructs of increasing length were placed under the control of the native *S. pyogenes* promoter, P_*native*_, or P_*tet*._ Efficiency of *lidA* targeting by crRNAs encoded by the terminal spacers of each array was assessed by monitoring LidA protein levels in Lp02(*dcas9*) and Lp02 by immunoblot. ICDH serves as a loading control. (C) A P_*tet*_*-*MC construct, MC-I, capable of expressing ten unique crRNAs was designed. crRNAs encoded by *S1, S3, S5, S6, S8*, or *S10* target genes in which antibodies directed against their encoded gene products are available. Bacteria pellets were collected from axenic cultures containing MC-I or the empty vector (+40 ng/mL aTC) and multiple immunoblots were run to accommodate the range of targets. ICDH serves as a loading control. Replicates are given in Figure S2.

To examine the ability of our multiplex CRISPRi platform to target a variety of genes simultaneously, we first made a synthetic MC array, called MC-I, composed of ten spacers (10-plex) under the regulation of P_*tet*_. The four spacers in position *S1, S3, S6*, and *S10* encoded crRNAs to target the effector-encoding genes *lidA, sidM, vipD*, and *ravN* (*lpg1111*), while the two spacers at position *S5* and *S8* target *dotD* (*lpg2674*) and *dotO* (*lpg0456*), two components of the type IV secretion system (T4SS) of *L. pneumophila*. Construction of MC constructs is described in the Methods section and depicted in Figure S1B. After growth of Lp02(*dcas9*) containing MC-I in axenic culture, we found that, indeed, the protein levels of all six targets were notably decreased based on immunoblot analysis as compared to a control strain bearing an empty vector (with tracrRNA but no repeat/spacer array) (Figure 2C and Figure S2). This initial result showed that it is possible to simultaneously silence several unique genes in *L. pneumophila* using an array of repeats and spacers under the control of a strong promoter.

### Spacer position influences gene silencing efficiency in P_*tet*_*-*MC arrays

Encouraged by the finding that our MC-I 10-plex construct had silenced multiple target genes, we determined how silencing efficiency varied dependent on the spacer position within the CRISPR array. To that end, we designed five additional 10-plex constructs, MC-II through MC-VI (Figure 3A), in which the aforementioned spacers encoding crRNA^lidA^, crRNA^vipD^, crRNA^sidM^, crRNA^dotO^, crRNA^dotD^, and crRNA^ravN^ were gradually shifted from proximal positions into more distal positions within the array, occupying either odd- or even-numbered positions. crRNA^lpg2793^ and crRNA^lpg0208^ were also encoded by spacers at various positions as these crRNAs had emerged in preliminary studies to silence very efficiently or not at all, respectively.

**Figure 3:**
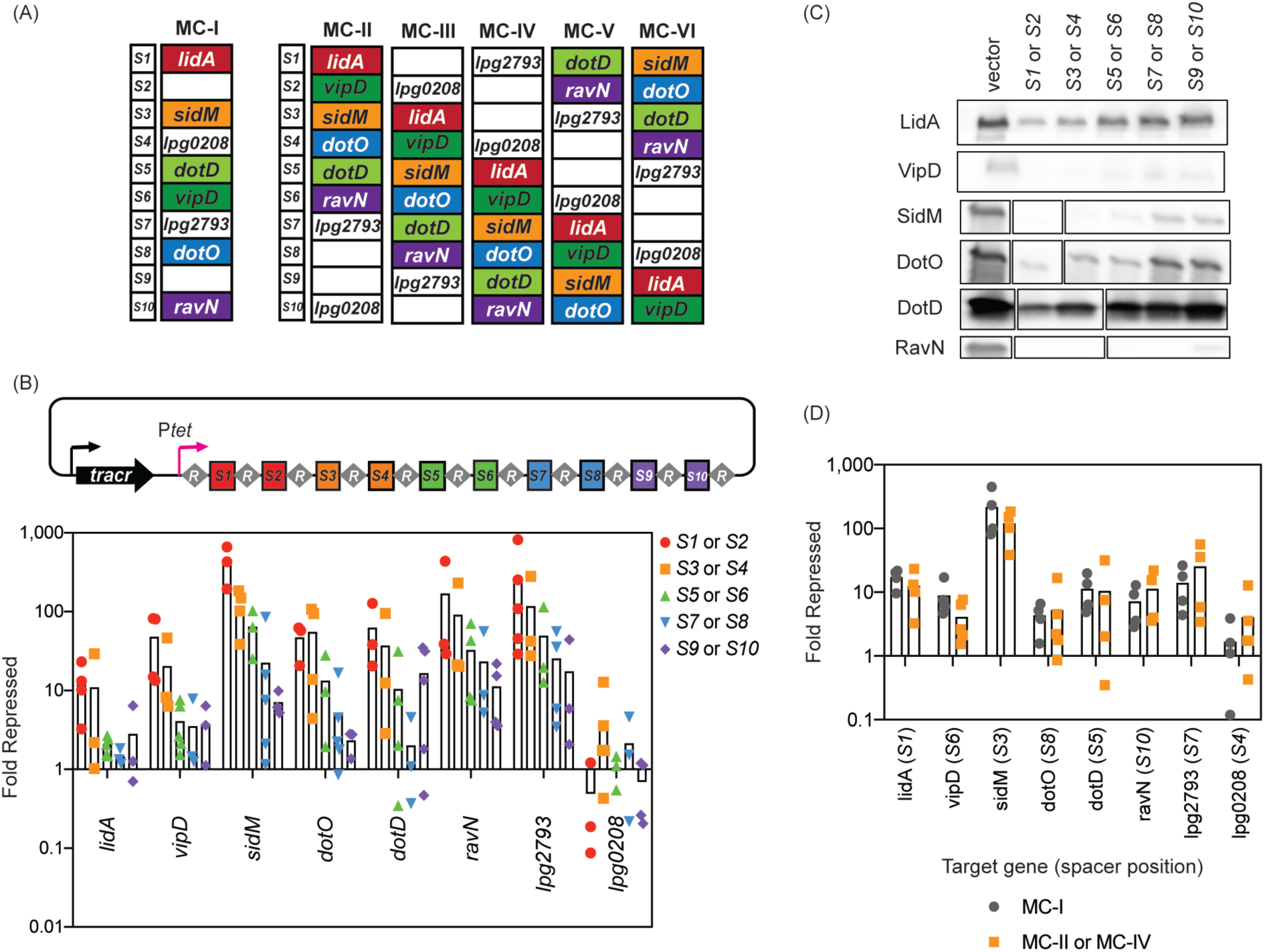
The effect of spacer position in gene silencing by P_*tet*_*-*MC arrays. (A) Additional P_*tet*_*-*MC constructs, MC-II through MC-VI, were made to test the effect of spacer position and environment on gene silencing. Each spacer is present in either position *S1, S3, S5, S7*, and *S9* or *S2, S4, S6, S8*, and *S10*. (B) RNA was extracted from Lp02(*dcas9*) bearing either the empty vector or an MC construct after axenic culture (+40 ng/mL aTC). qPCR was used to measure mRNA levels of target genes. Fold repression was determined using ΔΔC_T_. Bars indicate the mean of multiple replicates shown as individual data points. A value of 1 indicates that the mRNA level was the same in both strains. (C) Immunoblot analyses were performed on pellets collected from the same cultures. The blot shown is a representative of three replicates, with bands rearranged corresponding to spacer position. Original immunoblots and ICDH loading controls are shown in Figure S3. (D) RNA was extracted from Lp02(*dcas9*) bearing either the vector or the MC-I construct after axenic culture (+40 ng/mL aTC). qPCR was used to measure mRNA levels of target genes. Fold repression was determined using ΔΔC_T_ and plotted in comparison to data first shown in (B).

Efficiency of gene silencing by MC-II through VI after two days of induction during axenic growth was assessed for all targets at both the mRNA level and protein level and compared to that of the Lp02(*dcas9*) control strain containing the empty vector. qPCR analyses indeed revealed a polarity in gene silencing along the length of the repeat/spacer arrays, as has been noted for other CRISPRi platforms before (21, 33), with spacers in the proximal and distal positions causing the most- or least-efficient gene silencing, respectively (Figure 3B). Some crRNAs did silence their target genes very well (crRNA^sidM^, crRNA^ravN^, and crRNA^lpg2793^) even when encoded from distal spacer positions. In fact, *ravN* was as efficiently silenced by crRNA^ravN^ encoded by the *S10* spacer (MC-IV) as *lidA* was by crRNA^lidA^ encoded by the *S1* spacer (Figure 3B). crRNA^lpg2793^, which showed continuously high levels of repression, experienced greater than an order of magnitude repression even with the crRNA encoded by spacer *S10*. Conversely, crRNA^lpg0208^ did not work well regardless of the encoding spacer position. Notably, all genes except *lpg0208* experienced at least an order of magnitude repression when targeted by crRNAs encoded from spacers at positions as distal as *S4*, and five of them showed at least an order of magnitude silencing up to spacer position *S6*. Immunoblot analysis of the six targets for which antibodies were available confirmed that protein levels correlated very well with the mRNA abundance (Figure 3C and Figure S3), with depletion being near or below the detection limit.

### Gene silencing efficiency by CRISPRi is independent of spacer environment

crRNAs within the various MCs (MC-II through MC-VI), although transcribed from spacers in different positions, were always flanked by the same pair of neighboring spacers. For example, the spacer encoding the less efficient crRNA^lidA^ was always positioned at the 3’ and 5’ ends by spacers encoding crRNA^lpg0208^ and crRNA^vipD^, respectively. Those neighboring spacers may have inadvertently affected transcription or processing of crRNA^lidA^. To test if the spacer environment had an effect on crRNA production and therefore gene silencing efficiency, which has been proposed in the past (34), we compared target gene repression by crRNAs that were encoded by spacers at identical positions but surrounded by different spacers. crRNA^lidA^, crRNA^sidM^, and crRNA^dotD^ are encoded from the same position in MC-I and MC-II (spacers *S1, S3*, and *S5*), while crRNA^lpg0208^, crRNA^vipD^, crRNA^dotO^, and crRNA^ravN^ originate from position *S4, S6, S8*, and *S10* in both MC-I and MC-IV, yet they are flanked by entirely different sets of spacers (Figure 3A). Upon expression in Lp02(*dcas9*), crRNAs encoded by MC-II or MC-IV, despite being encoded from different spacer environments, consistently achieved the same level of gene repression as those encoded by MC-I (Figure 3D). Together, these results demonstrate that spacer position along the array has a much greater impact on multiplex CRISPRi-mediated gene silencing than the spacer environment.

### *boxA* elements reduce polarity of spacer transcription to promote maximum gene silencing

While the addition of the strong P_*tet*_ promoter did enhance gene silencing by distal spacers (Figure 2B), gene silencing polarity within the MC arrays was still not fully overcome (Figure 3), indicating that additional factors likely influenced crRNA production. *BoxA* elements are often found upstream of long non-coding RNAs, like the 16S RNA (35), where they are bound by the Nus complex to prevent Rho-dependent transcription termination (36-38). *Stringer et. al* recently discovered *boxA* elements to be evolutionarily conserved upstream of naturally occurring CRISPR arrays where they appear to promote expression of downstream spacers within long arrays (39). To investigate whether Rho-dependent transcriptional termination had prevented our CRISPRi platform from achieving maximum gene silencing from distal spacer positions, knowing that *L. pneumophila* does encode at least one member of the Nus complex (39), we explored if incorporation of *boxA* elements could overcome this limitation. We compared the *E. coli boxA* consensus sequence with that of the *Coxiella* 16S *boxA* and the *boxA* found upstream of the naturally occurring *L. pneumophila sp. Lens* CRISPR array (Figure 4A). Interestingly, a *boxA* element was not readily identified upstream of the *L. pneumophila* Philadelphia-1 16S RNA encoding region. Thus, we mutated the leader sequence of the repeat/spacer array of MC-II through MC-VI to include the 11 base pair *L. pneumophila sp. Lens boxA* sequence (5’-GTTCTTTAAAA-3’), and the five flanking base pairs on either side, at both a distance of -58 (*boxA*(−58)) and -90 (*boxA*(−90)) base pairs upstream of the first repeat while maintaining the overall length of the leader sequence (see Figure S4 for sequence detail).

**Figure 4:**
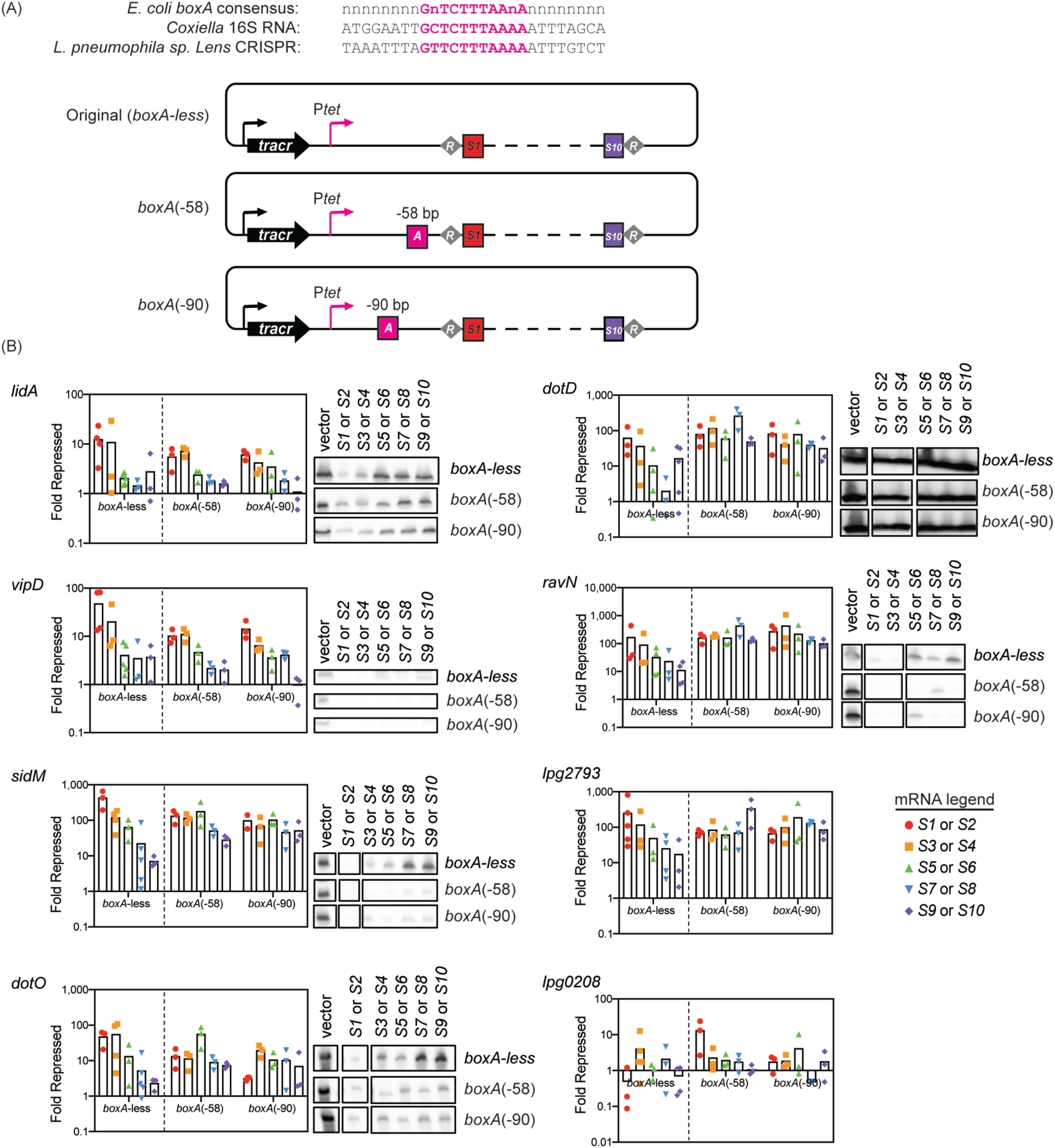
The effect of *boxA* elements on gene silencing by P_*tet*_*-*MC arrays. (A) *boxA* elements identified upstream of the *Coxiella* 16S RNA encoding sequence and the *L. pneumophila sp. Lens* CRISPR array were compared to that of the *E. coli boxA* consensus. The *boxA* motif from the *L. pneumophila sp. Lens* CRISPR array was added to the leader sequence of the P_*tet*_*-*MC constructs from Figure 3 at either -58 (*boxA*(−58)) or -90 (*boxA*(−90)) base pairs upstream of the repeat/spacer array. *boxA* is depicted as a pink square. (B) RNA extracted from axenic cultures of *boxA* construct-bearing Lp02(*dcas9*) was compared to that of the empty vector-bearing Lp02(*dcas9*) by qPCR, as in Figure 3 (+40 ng/mL aTC). *boxA*-less data from Figure 3 is shown again for comparison to the new *boxA*(−58) and *boxA*(−90) data. Immunoblot analyses were performed on pellets collected from *boxA*-less, *boxA*(−58) and *boxA*(−90) grown side-by-side. The bands were reordered to correspond to spacer position. Original immunoblots and ICDH loading controls are shown in Figure S5.

Upon providing Lp02(*dcas9*) with these *boxA* constructs, mRNA and protein analysis (Figure 4B) revealed that this *boxA* element greatly resolved gene silencing polarity for the majority of the target genes tested. The presence of *boxA* in either the -58 or -90 position maximized silencing of *sidM, dotO, dotD, ravN*, and *lpg2793* by crRNAs encoded from all spacer positions, even *S9* or *S10*, presumably by allowing transcription of the repeat/spacer array to persist throughout the length of the array. *sidM, dotD, ravN*, and *lpg2793* displayed nearly two orders of magnitude gene repression from crRNAs encoded from all spacer positions, a vast improvement over the diminishing repression observed by *boxA*-less constructs. Silencing of *lidA, lpg0208*, and *vipD* was not altered by the addition of *boxA* suggesting that other factors such as a strong promoter or contending transcriptional activators were contributing to the expression of the target genes and were possibly inhibiting silencing efficiency. In conclusion, the processivity of transcription of long repeat/spacer arrays is the major driving factor of achieving multiplex gene silencing even from distal spacer positions.

### CRISPRi is functional during *L. pneumophila* intracellular growth

While CRISPRi has been performed in a variety of bacteria during growth in axenic culture, gene silencing had yet to be accomplished in any pathogen during infection. We confirmed that our platform was functional in *L. pneumophila* undergoing intracellular growth and that, upon repression of specific targets, we could reproduce previously reported intracellular growth phenotypes.

*L. pneumophila* requires the Dot/Icm T4SS to deliver effector proteins into the host cell in order to establish a replication vacuole. Strains that bear loss of function mutations in the genes encoding T4SS subunits are unable to establish this specialized vacuole and fail to replicate (40). The ATPase DotO and the outer membrane protein DotD are components critical for the function of the T4SS transporter, and *L. pneumophila* mutants with disruptions in their encoding genes (*lpg0456* and *lpg2674*, respectively) are attenuated for virulence (41-43). Plasmids encoding crRNAs directed against *dotO* (crRNA^dotO^) or *dotD* (crRNA^dotD^) were individually introduced to Lp02(*dcas9*) and, after 48 hours of induction, decreased protein levels of DotO and DotD were verified by immunoblot analysis (Figure 5A). It is important to note, and well established (41, 42), that mutations disabling the T4SS have no influence on the general fitness of *L. pneumophila* outside the context of the host. Human-derived U937 macrophages (44, 45) were then challenged with these crRNA-expressing strains, and bacterial growth was monitored over a period of 72 hours (Figure 5A). While the Lp02(*dcas9*) strain containing the empty plasmid grew several orders of magnitude, the CRISPRi strains depleted of either DotO or DotD were impaired for growth at a level comparable to that of the avirulent mutant strain Lp03 which bears a chromosomal mutation in the *dotA* gene that encodes another critical subunit of the T4SS (27).

**Figure 5:**
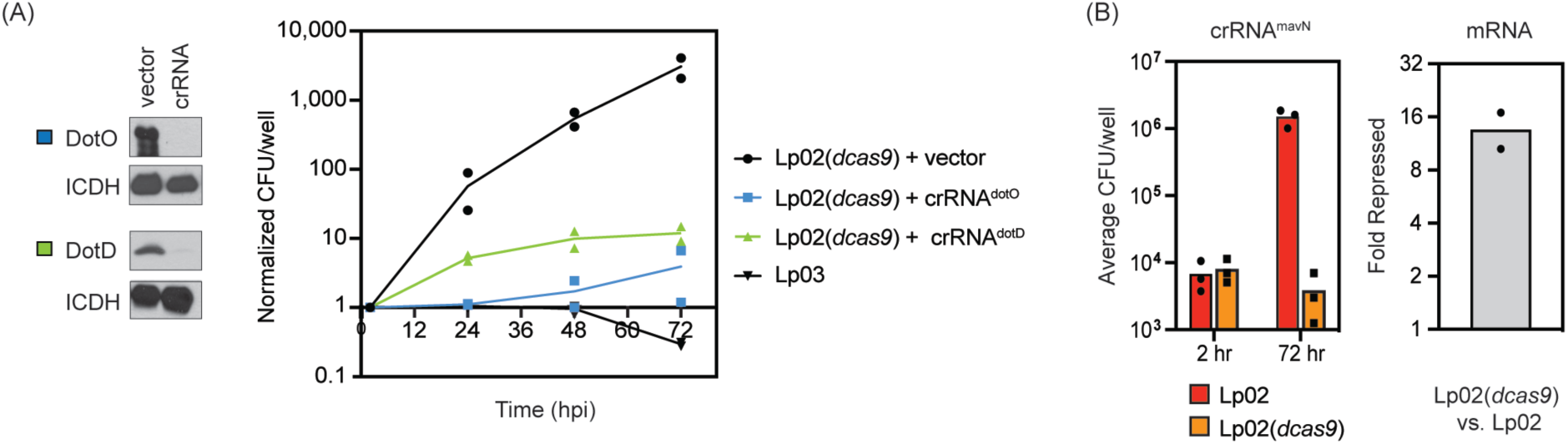
*L. pneumophila* CRISPRi is functional during intracellular growth. (A) Immunoblots confirm decreased levels of DotO and DotD in Lp02(*dcas9*) strains expressing crRNA^dotO^ or crRNA^dotD^. ICDH serves as a loading control. Growth of these strains in U937 macrophages was compared to that of Lp02(*dcas9*) bearing an empty vector and the avirulent strain Lp03, with a chromosomal mutation in *dotA*. Growth is plotted as CFU/well normalized to the CFU at 2 hours post infection (hpi). (B) Growth of Lp02 and Lp02(*dcas9*) expressing crRNA^mavN^ was assayed in U937 macrophages. Growth is given as the average of CFU/well of three experiments at 2 and 72 hpi. mRNA levels of *mavN* in Lp02(*dcas9*) *vs*. Lp02 were measured by qPCR. Fold repression was determined using ΔΔC_T_.

In agreement with this result, our efforts to target and repress an effector-encoding gene also proved successful. MavN is a multi-spanning membrane protein from *L. pneumophila* that is essential for delivering metal ions across the LCV membrane into the vacuolar lumen (46, 47). Deletion of *mavN* (*lpg2815*) from the *L. pneumophila* chromosome results in a complete growth defect in A/J mouse macrophages (46). Upon targeting *mavN* with crRNA^mavN^, we observed a similarly dramatic growth defect in U937 macrophages in the Lp02(*dcas9*) strain, while the Lp02 control strain grew robustly over the same time period (Figure 5B). In the absence of an antibody specific to MavN, mRNA levels were analyzed by qPCR and confirmed *mavN* to be repressed more than ten-fold in the Lp02(*dcas9*) strain relative to the Lp02 control strain. These data indicate that CRISPRi is functional in *L. pneumophila* both during growth in axenic media and during a multi-day infection experiment.

### Multiplex CRISPRi reveals gene dosage effects during intracellular growth

At last, we explored if intracellular growth of *L. pneumophila* can be manipulated through multiplex CRISPRi. We created *dotO*-specific P_*tet*_*-*MC constructs (called MC-II-dotO to MC-VI-dotO; Figure 6A) that mimicked P_*tet*_*-*MC-II through MC-VI, except that all crRNA-encoding spacers besides the crRNA^dotO^-encoding spacer were scrambled, rendering them incapable of base pairing with their original targets. Notably, the base pair count and composition (“agtc” content) of the scrambled spacers were maintained, and all repeat sequences remained intact. As a result, these multiplex arrays should encode only crRNA^dotO^ as functional crRNA from spacer positions *S2, S4, S6, S8*, or *S10* for direct comparison to the single crRNA^dotO^-encoding CRISPRi construct (from Figure 5). Furthermore, we created a control construct, MC-VII-dotO, in which the *dotO*-targeting spacer in position *S2* was scrambled to provide evidence that any observed phenotypes were a direct result of silencing *dotO*.

**Figure 6:**
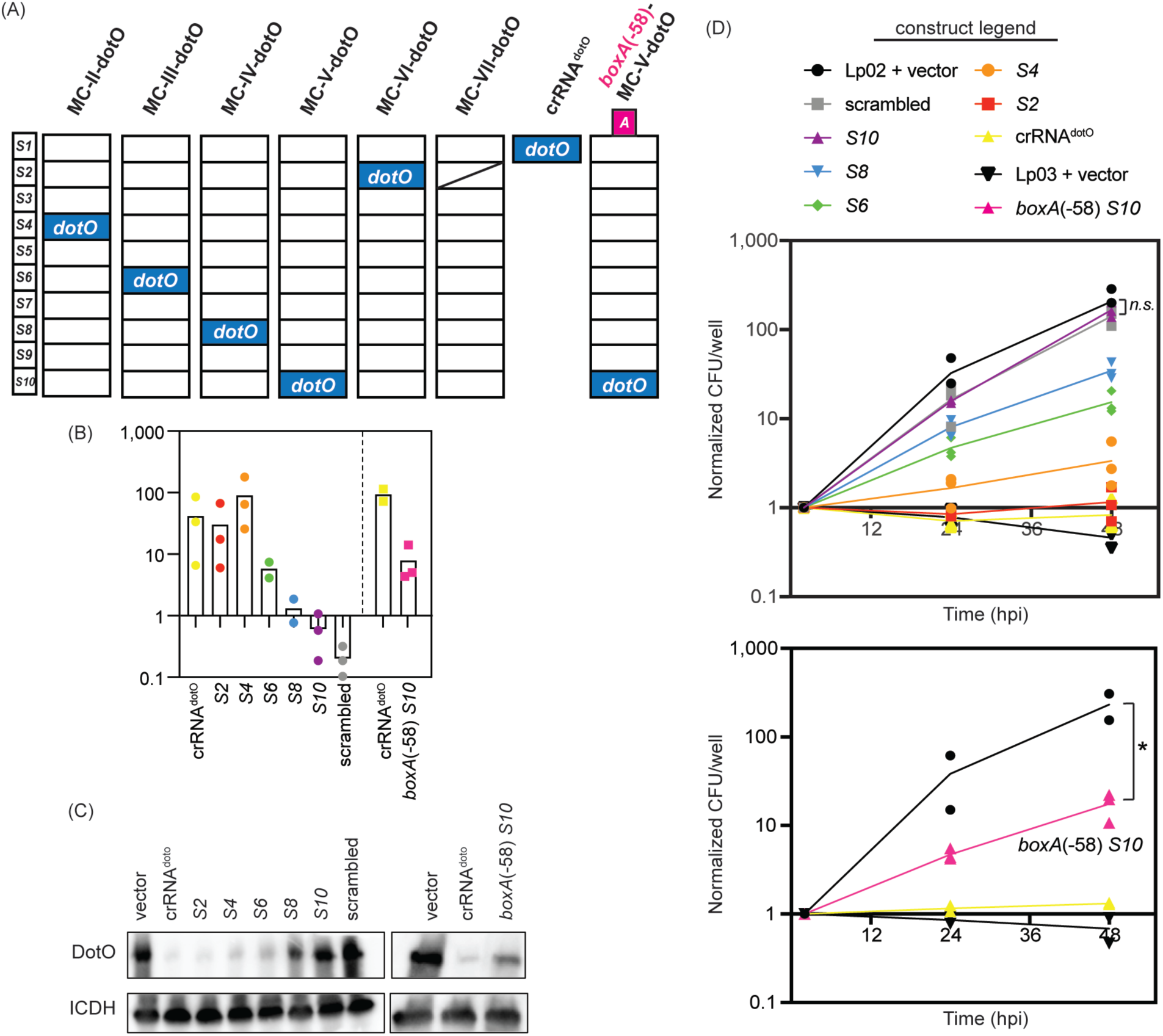
Multiplex CRISPRi is functional in *L. pneumophila* during host infection. (A) *boxA*-less, *dotO*-targeting P_*tet*_*-*MC constructs, MC-II-dotO, MC-III-dotO, MC-IV-dotO, MC-V-dotO, and MC-VI-dotO, were constructed such that the sequences of all spacers, other than *dotO*, were scrambled. The *dotO* spacer can be found in the same position as in the original MC-II through MC-VI constructs. MC-VII-dotO is identical to MC-VI-dotO except that the *dotO* spacer is also scrambled. crRNA^dotO^ is the same as in Figure 5. Later, a *boxA* sequence was added in the -58 position to MC-V-dotO to create *boxA*(−58) *S10*. (B) RNA extracted from axenic cultures of Lp02(*dcas9*) bearing these constructs was compared to that of Lp02(*dcas9*) bearing the empty vector by qPCR, as in Figure 3 (+40 ng/mL aTC). (C) Immunoblot analyses were performed on pellets collected from the same cultures and samples were run according to *dotO* spacer position. Immunoblot replicates are found in Figure S6. ICDH serves as a loading control. (D) aTC-induced Lp02(*dcas9*) bearing *boxA-*less and *boxA*(−58) constructs were used to infect U937 macrophages. Growth of these strains and Lp02(*dcas9*) and Lp03 bearing the empty vector was monitored over 48 hours post infection (hpi). Colony forming unit (CFU) counts were normalized to the count at two hours post infection. Normalized counts for each experiment at each timepoint are shown. *n*.*s*. = not significant (*p=0*.*44, n=*3), \**p=*0.033, *n=*2 or 3, one-unpaired *t*-test.

We examined *dotO* repression by the new constructs in axenic culture to establish the degree of *dotO* silencing efficiency. qPCR revealed that *dotO* silencing by MC-VI-dotO and MC-II-dotO constructs with crRNA^dotO^-encoding spacers in the proximal *S2* and *S4* positions, respectively, was robust and comparable to that of the single crRNA^dotO^-encoding CRISPR construct (Figure 6B). *dotO* silencing efficiency by the MC-III-dotO to MC-V-dotO constructs gradually decreased with increasing distal shift of the crRNA^dotO^-encoding spacer in these arrays. As expected, the scrambled *dotO* spacer in MC-VII-dotO did not promote gene silencing, showing that the effects of MC-II-dotO to MC-VI-dotO was specific to crRNA^dotO^. Again, immunoblot analyses of cell pellets collected from these axenic *L. pneumophila* cultures showed depletion of DotO to the same extent as would be predicted from the mRNA abundance data (Figure 6C and Figure S6).

We then challenged U937 macrophages with these *boxA*-less crRNA^dotO^-expressing strains and compared their growth over a period of 48 hours to that of Lp02(*dcas9*) with the empty vector, which served as a measure of maximum growth, and Lp03, which served as a measure of maximum attenuation (Figure 6D, top). Strikingly, the severity of the growth defect precisely mirrored the reduction in *dotO* mRNA levels caused by these constructs, with the most severe defect caused by crRNA expression from the *S2* spacer (MC-VI-dotO construct) and lesser defects observed with the distal shift of the *dotO* spacer in MC-II-dotO to MC-V-dotO. The scrambled *dotO* spacer of MC-VII-dotO was unable to silence *dotO* and allowed proficient *L. pneumophila* growth comparable to the Lp02 control strain. Thus, the polarity of spacer expression and the corresponding decay of target repression represents a titrated gene silencing platform.

Since silencing was least efficient for the MC-V-dotO construct (*S10*, purple triangle in Figure 6D, top), we added a *boxA*(−58) motif to the leader sequence of the array and re-examined the effect of this *boxA*(−58)-MC-V-dotO construct on *L. pneumophila* intracellular growth. Remarkably, we now observed a significant attenuation of growth in U937 macrophages over 48 hours, achieving over an order of magnitude replication defect compared to Lp02(*dcas9*) bearing the empty vector (Figure 6D bottom, *p=*0.033, one unpaired *t-*test). In fact, with the *boxA*(−58) motif present, the growth defect observed by expressing the crRNA from the *S10* spacer was now comparable to that of expression from the *S6* spacer in a *boxA*-less construct (green diamond in Figure 6D, top). mRNA and protein analyses confirmed that *dotO* silencing by the *boxA*(−58)-MC-V-dotO construct mirrored that of the *dotO* silencing in the *boxA*-less MC-III-dotO construct (Figures 6B and 6C). Together, these data demonstrate that multiplex gene silencing by long CRISPR arrays can be efficiently boosted even within infection models simply by positioning a *boxA* element in the leader region where it mediates anti-termination.

## Discussion

In this study, we have established a multiplex CRISPRi platform in the pathogen *L. pneumophila* and provide proof-of-concept for this novel platform to be usable not only during growth in axenic media but also during macrophage infection where it reproduced known intracellular growth phenotypes (Figure 6). Importantly, by placing the crRNA-encoding spacer in positions further downstream within the array, the degree of gene silencing was titratable (Figure 3). In contrast, when combined with a *boxA* anti-transcription termination element, our 10-plex CRISPR array had the potential to silence up to ten unique genes simultaneously (Figure 4) making it a powerful tool to study even synergistic genetic interactions.

Our methodical approach to analyze the potential of multiplex CRISPRi not only allowed us to generate a powerful platform capable of silencing up to ten unique genes, a platform that may prove invaluable for studying the more than 300 *L. pneumophila* effectors in the future, but it also revealed additional surprises that could be applied to future gene interrogations. For example, some of our MC arrays encoded not just one but two crRNAs (called *lidA1* and *lidA2*) towards the same target gene (Figure 2B), resulting in LidA to be efficiently depleted even though the encoding spacers were located in most distal spacer positions (*S9* and *S10*) within the array. Notably, when only the crRNA encoded from the *lidA1* spacer was used in subsequent MC constructs (Figure 3), silencing was not nearly as efficient, suggesting that the combined action of both spacers in P_*tet*_-MC-9/10 promoted maximum silencing. While only a single example, future users of this technology could experiment by adding more than one spacer for a given gene to enhance gene silencing. crRNAs encoded by spacers *lidA1* and *lidA2* target different regions of *lidA*, but one could hypothesize repeating the same spacer sequence could also increase gene silencing of a target by producing more of the crRNA. The malleability of our MC scaffold allows users who may not need to target ten unique genes to apply these tactics to achieve maximum silencing levels of fewer gene targets.

Our platform also provides two different strategies for titratable gene silencing. First, since both *dcas9* and the CRISPR array are under the regulation of the P_tet_ promoter, simply altering the amount of aTC inducer presumably changes gene silencing by controlling both the quantity of dCas9 and the abundance of the crRNA (Figure 1B). While regulating crRNA expression with a variety of inducers (*e*.*g*. aTC, arabinose, xylose, and IPTG) is common place, gene silencing in this way has been argued to be noisy (48). Second, titrated gene silencing was also accomplished when P_*tet*_-MC constructs without a *boxA* motif were used that were prone to transcription termination (Figure 3 and 6). DotO constructs provided a clear example of how gene silencing through placing spacers at ever distant positions within the array can lead to titrated effects to the biological system (Figure 6D). Recent efforts have been made to tailor the degree of crRNA-targeting of a gene by creating numerous variations of the spacer sequence away from the perfect match (48, 49). We imagine shifting the ideal spacer downward in the array would be a much simpler feat.

Using CRISPRi in *L. pneumophila* allowed for the systematic investigation of not only repeat/spacer arrays as a gene targeting platform, but also provided additional insight into how CRISPR arrays might function in nature. We show that with the native *S. pyogenes* promoter alone, only the most proximal spacers are transcribed (Figure 2B). It is known that spacers acquired from most recent infections are inserted at the first position in the array (50). Our data reiterate that this is an advantageous strategy as the earliest spacers in the array are the most likely to be transcribed, and therefore, will produce the greatest quantity of protective crRNAs for an ongoing viral attack. Several bacterial CRISPR systems appear to have overcome the limitation of promoter-driven transcription processivity by inhibiting transcription termination through acquiring Nus factor-binding *boxA* sequences upstream of their arrays (39). Adding the *boxA* sequence to the synthetic array also presumably increased transcription processivity of our synthetic constructs, as genes targeted by crRNAs encoded by spacers in more distal positions were now efficiently silenced (Figure 4). Still, as naturally occurring CRISPR arrays can stretch to the hundreds of spacers (32), there could be additional unknown factors that regulate spacer transcription that could one day be applied to a CRISPR-based tool.

Ultimately, our *dotO*-silencing experiments during host infection provided evidence that this multiplex CRISPRi technology is ready to be applied to biological investigations, especially in the study of *L. pneumophila* pathogenesis. As touched on above, *dotO* expression appears to be exquisitely tailored towards promoting maximum infection, as any deviation of expression lead to growth attenuation in macrophages (Figure 6). In what could be a nutrient limited environment, it seems *L. pneumophila* expresses just enough DotO in nature to achieve virulence through T4SS assembly and secretion of effectors, without being wasteful by producing an excess amount of this protein.

The *Legionella* community has long thought that the multitude of *L. pneumophila* effectors combined with the lack of growth phenotypes observed upon creating single deletion strains supports the existence of redundancy and synergy amongst the effectors (51). Looking forward, the multiplex CRISPRi approach developed here holds the promise of one day probing functional overlap amongst the hundreds of *L. pneumophila* effectors. Not only can genes be silenced in bulk groups, but the mobility of our single plasmid based CRISPRi platform allows for easy transfer of MC constructs into a variety of *Legionella* mutant strain backgrounds to directly assess redundancy, presuming they have been equipped with a copy of *dcas9*. These analyses are not limited to macrophage infections, but any number of hosts. In this capacity, multiplex CRISPRi serves as an initial discovery tool that provides guidance on which proteins to focus on during follow-up analyses. When adapted for use in other microbial pathogens, the multiplex CRISPRi technology developed here has the potential to promote understanding of their biology, and possibly even foster the discovery of prospective drug targets.

## Materials and Methods

### Construction of *dcas9*-containing strains

*dcas9* was added to the chromosome of *Legionella pneumophila* Philadelphia-1 Lp02 (*thyA hsdR rpsL*) through allelic exchange to make MML109 (Lp02(*dcas9*). The tet^R^-*dcas9* segment of pdCas9-bacteria (Addgene #44249) was amplified with BKMP108-109 and introduced into pDonorP4r-P3r (Invitrogen) to generate pMME1080. The N- and C-terminus of *thyA* (*lpg2868*) were amplified by BKMP94-95 and BKMP98-99 primers and introduced into pDonorP1-P4 and pDonorP3-P2 (Invitrogen) to generate pMME1084 and pMME1094, respectively. These three donor vectors were introduced into pNPTS138_Cm-DEST (pMME1020, constructed by placing the Gateway DEST sequence on pNPTS138_Cm (Addgene #41891) at HindIII and SpeI) by a Gateway LR reaction (Invitrogen) to generate pMME1115 (N-terminus *thyA*::tet^R^-*dcas9* cassette::C-terminus *thyA*). pMME1115 was introduced into Lp02 by electroporation and strains containing the plasmid were selected for on CYET-Chloramphenicol (Cm) plates (CYET plates described previously (52)). Cm-resistant colonies were patched on CYET plates containing 5% sucrose to remove the plasmid backbone, leaving behind the N-terminus *thyA*::tet^R^-*dcas9* cassette::C-terminus *thyA* by homologous recombination. *dcas9* incorporation into *Legionella* chromosome was confirmed by immunoblot analyses and whole genome sequencing. Primer sequences are listed in Table S3.

### Construction of single CRISPRi constructs and preliminary multiplex CRISPRi arrays

crRNA-expressing constructs were built using Invitrogen Gateway-compatible plasmids as depicted in Figure S1A. The BsaI-CRISPR segment of pCRISPR (Addgene #42875) was amplified with BKMP185-186 and introduced into pDonorP5-P2 (Invitrogen) to generate pMME1540. Specific single crRNA-encoding spacer sequences were designed as explained in Figure 1 and added to the BsaI-CRISPR segment of pMME1540 as previously described for pCRISPR (29). To make preliminary multiplex CRISPRi constructs as shown in Figure 2B, 21 crRNA-encoding spacers separated by repeats were synthesized by GenScript and provided on a pUC57 vector (pMME1170). CRISPR backbones containing four (MC-3/4), eight (MC-7/8), or ten (MC-9/10) crRNA-encoding spacers were amplified from this plasmid using forward primers crsidF_F, crsidD_F, crsetA_F, respectively, and reverse primer crlidA2_R. PCR products were BsaI treated and ligated into pMME1540 (P_*native*_) or pMME1748 (P_*tet*_) as above.

Next, the *Streptococcus pyogenes* tracrRNA-encoding sequence from pCas9 (Addgene #42876) was amplified with BKMP45-46 and introduced into pDonorP1-P5 (Invitrogen) to generate pMME985. The crRNA donor plasmids, for both the single CRISPRi constructs and the preliminary multiplex CRISPRi arrays, and pMME985 were introduced into pMME977 by the Gateway LR reaction to generate the final crRNA-encoding plasmids. Final plasmids were introduced to Lp02(−*dcas9*) or Lp02(*dcas9*) by electroporation and the strains containing the plasmid were selected for on CYE media. All final strains and crRNA-encoding spacer sequences are listed in Table S2. Primer sequences are listed in Table S3.

### Construction of P_*tet*_-MC multiplex CRISPRi constructs

Multiplex crRNA-expressing constructs were built from GenScript-synthesized plasmids as depicted in Figure S1B and the nucleotide scaffold for P_*tet*_-MC given in Figure S7A. MC-I through MC-VI and MC-II-dotO through MC-VII-dotO array sequences were provided on a pUC57 vector (Table S1). Next, the MC array within pUC57 was amplified using Gateway5-Ptet and Gateway2-T1term and introduced into pDonorP5-P2 by the Gateway BP reaction to generate donor plasmids. The MC backbone vector, containing simply the leader sequence of the multiplex CRISPRi array, was amplified from a GenScript plasmid using Gateway5-leader and Gateway2-leader and similarly introduced into pDonorP5-P2. These donor plasmids and pMME985 were introduced into pMME977 by the Gateway LR reaction to generate the multiplex CRISPRi constructs listed in Table S2. These plasmids were then introduced to Lp02(*dcas9*) by electroporation and the strains containing the plasmid were selected for on CYE plates without thymidine. All final strains and crRNA-encoding spacer sequences are listed in Table S2. Primer sequences are listed in Table S3.

### Addition of *boxA* to P_*tet*_-MC multiplex CRISPRi constructs

The *boxA* sequence from *L. pneumophila sp. Lens* was introduced to the leader region of the MC-II through MC-VI and MC-V-dotO arrays in our donor plasmids by quickchange PCR with Pfu Turbo Polymerase (Agilent #600250-52). *boxA*(−58) was added using primers BoxA_srtF/R and *boxA*(−90) was added using primers BoxA_ farF/R. Then sequence-verified donor plasmids and pMME985 were introduced into pMME977 by the Gateway LR reaction as before. Again, these plasmids were introduced to Lp02(*dcas9*) by electroporation and the strains containing the plasmid were selected for on CYE plates. The nucleotide scaffold for *boxA*(−58) P_*tet*_-MC is given in Figure S7B. All final strains and crRNA-encoding spacer sequences are listed in Table S2. Primer sequences are listed in Table S3.

### Axenic growth of *L. pneumophila* CRISPRi strains

For experiments with single CRISPRi constructs (Figures 1 and 5), *L. pneumophila* were grown overnight in AYE (10 g ACES, 10 g yeast extract per liter, pH 6.9 with 0.4 mg/ml cysteine and 0.135 mg/ml ferric nitrate) under inducing conditions (by adding either 20 ng/mL or 40 ng/mL anhydrous tetracycline (aTC, Clontech #631310)). For all other experiments, *L. pneumophila* cultures were grown overnight in AYE under non-inducing conditions (−aTC). On the second day, cultures were sub-cultured twice (AM and PM, ∼6-7 hours apart) to OD_600_ 0.2-0.3 with 2-3 mL fresh AYE containing 40 ng/mL aTC. On the third day, cultures that had reached OD_600_ 3-5 (post-exponential growth) were collected for mRNA analyses, immunoblot analyses, and/or use in host cell infections.

### Immunoblot assays and antibodies

Immunoblot assays were performed on bacteria pellets resuspended in SDS sample buffer to either 1×10^8^ CFU or OD_600_ =10. SDS-PAGE gels were run on a Protein III system and transfers were performed with the Trans-Blot Turbo Transfer System (BioRad). Protein bands were detected using a primary antibody followed by an HRP-conjugated secondary antibody (rabbit, Life technologies #G21234, or mouse, Invitrogen #G21040) and visualized via chemiluminescence using Clarity Western ECL Substrate and the ChemiDoc MP Imaging System (BioRad). Primary antibodies directed against SidM (53), VipD (54), RavN (55), and LidA (56) were described before. Primary antibody against ICDH was a kind gift of Abraham (Linc) Sonenshein (Tufts University School of Medicine). Primary antibodies against DotD and DotO were a kind gift of Joseph Vogel (Washington University in St. Louis). Anti-Cas9 monoclonal antibody was purchased from Active motif (Catalog #61577).

### RNA extraction and qPCR

Bacterial RNA extraction was performed on bacteria pellets using the Trizol Max Bacterial RNA Isolation Kit (Invitrogen #16096040). Contaminating DNA was removed using the Turbo DNA-free Kit (Invitrogen #AM1907) and RNA was converted to cDNA using the High-Capacity cDNA Reverse Transcription Kit (Applied Biosystems #4368814). qPCR was performed using the SYBR Green Master Mix (Applied Biosystems #4367659) on a StepOnePlus Real-Time PCR System (Applied Biosystems) using comparative C_T_ and the standard 2-hour protocol. qPCR primers (found in Table S3) were designed using NCBI Primer-BLAST such that they amplified ∼100-150 bp of sequence near the 5’ end of the gene of interest and had a melting temperature between 57 °C and 63 °C. mRNA levels from different samples were normalized to the house-keeping gene (*rpsL*) levels and mRNA levels in CRISPRi strains were compared to that of -dCas9 strains or the empty vector control strain using the ΔΔC_T_ method (57) to determine fold repression.

### *L. pneumophila* intracellular growth assay in human derived U937 cells

U937 monocytes (ATCC CRL-1593.2) were maintained in DMEM + 10% FBS + glutamine. Three days prior to challenge with *L. pneumophila*, cells were plated on 24-well plates at 3×10^5^ cells/well with 0.1 µg/mL 12-O-tetradecanoylphorbol-13-acetate (TPA, Sigma-Aldrich #P1585) to promote differentiation. *L. pneumophila* strains were incubated under inducing conditions, as described above, and added to differentiated U937 macrophages in DMEM + 10% FBS + glutamine containing 40 ng/mL aTC at a multiplicity of infection (MOI) of 0.05. Plates were centrifuged for five minutes at 200 x g to increase cell-cell contact. After a 2-hour incubation, extracellular bacteria were removed by washing cells twice with DMEM +FBS + glutamine media containing 40 ng/mL aTC. Bacteria were collected 2, 24, 48, or 72 hours post infection (hpi). To extract the bacteria from the macrophages, digitonin (0.02% final concentration) was added to each well and incubated 10 min at 37 °C. Subsequently, lysate was collected, and each well was rinsed with dH_2_O to ensure collection of all bacteria. Bacterial samples were serially diluted and spotted on CYE plates to determine CFU. Results are given as the CFU relative to the CFU at 2 hpi.

### Statistical analysis of *L. pneumophila* growth

Statistical comparison of growth between MC-V-dotO and Lp02(*dcas9*) bearing the empty vector or *boxA*(−58)-MC-V-dotO and Lp02(*dcas9*) bearing the empty vector was carried out in GraphPad Prism 8 using the *t-*test analysis function. A one-unpaired *t-*test was performed without correction for multiple comparisons and without assuming a consistent SD. Changes in growth were considered significant if *p*<0.05.

## Acknowledgment

We thank members of the Machner laboratory for their critical reading of the manuscript, Dr. Joseph T. Wade (New York State Department of Health, Wadsworth Center) for advice and direction on *boxA* motifs, and Dr. Caroline Esnault (National Institutes of Health) for instruction on qPCR procedure. This work was funded by the Intramural Research Program of the National Institutes of Health, USA (Project Number: 1ZIAHD008893-07).

## Author Contributions

N.A.E., B.K., and M.P.M. contributed to experimental design; N.A.E. and B.K. performed experiments; N.A.E. and M.P.M. composed the manuscript.

## References

1. F. Hille et al., The Biology of CRISPR-Cas: Backward and Forward. Cell 172, 1239–1259 (2018).

2. M. Adli, The CRISPR tool kit for genome editing and beyond. Nat Commun 9, 1911 (2018).

3. J. R. Strich, D. S. Chertow, CRISPR-Cas Biology and Its Application to Infectious Diseases. J Clin Microbiol 57 (2019).

4. R. Barrangou et al., CRISPR provides acquired resistance against viruses in prokaryotes. Science 315, 1709–1712 (2007).

5. D. Bhaya, M. Davison, R. Barrangou, CRISPR-Cas systems in bacteria and archaea: versatile small RNAs for adaptive defense and regulation. Annu Rev Genet 45, 273–297 (2011).

6. E. Deltcheva et al., CRISPR RNA maturation by trans-encoded small RNA and host factor RNase III. Nature 471, 602–607 (2011).

7. S. J. Brouns et al., Small CRISPR RNAs guide antiviral defense in prokaryotes. Science 321, 960–964 (2008).

8. B. Wiedenheft, S. H. Sternberg, J. A. Doudna, RNA-guided genetic silencing systems in bacteria and archaea. Nature 482, 331–338 (2012).

9. Y. Zhu, S. E. Klompe, M. Vlot, J. van der Oost, R. H. J. Staals, Shooting the messenger: RNA-targetting CRISPR-Cas systems. Biosci Rep 38 (2018).

10. L. A. Marraffini, E. J. Sontheimer, Self versus non-self discrimination during CRISPR RNA-directed immunity. Nature 463, 568–571 (2010).

11. M. Jinek et al., A programmable dual-RNA-guided DNA endonuclease in adaptive bacterial immunity. Science 337, 816–821 (2012).

12. G. Gasiunas, R. Barrangou, P. Horvath, V. Siksnys, Cas9-crRNA ribonucleoprotein complex mediates specific DNA cleavage for adaptive immunity in bacteria. Proc Natl Acad Sci U S A 109, E2579–2586 (2012).

13. K. Chylinski, K. S. Makarova, E. Charpentier, E. V. Koonin, Classification and evolution of type II CRISPR-Cas systems. Nucleic Acids Res 42, 6091–6105 (2014).

14. L. S. Qi et al., Repurposing CRISPR as an RNA-guided platform for sequence-specific control of gene expression. Cell 152, 1173–1183 (2013).

15. D. Bikard et al., Programmable repression and activation of bacterial gene expression using an engineered CRISPR-Cas system. Nucleic Acids Res 41, 7429–7437 (2013).

16. M. H. Larson et al., CRISPR interference (CRISPRi) for sequence-specific control of gene expression. Nat Protoc 8, 2180–2196 (2013).

17. A. Vigouroux, D. Bikard, CRISPR Tools To Control Gene Expression in Bacteria. Microbiol Mol Biol Rev 84 (2020).

18. J. M. Peters et al., Enabling genetic analysis of diverse bacteria with Mobile-CRISPRi. Nat Microbiol 4, 244–250 (2019).

19. B. Adiego-Perez et al., Multiplex genome editing of microorganisms using CRISPR-Cas. FEMS Microbiol Lett 366 (2019).

20. J. M. Peters et al., A Comprehensive, CRISPR-based Functional Analysis of Essential Genes in Bacteria. Cell 165, 1493–1506 (2016).

21. D. Kaczmarzyk, I. Cengic, L. Yao, E. P. Hudson, Diversion of the long-chain acyl-ACP pool in Synechocystis to fatty alcohols through CRISPRi repression of the essential phosphate acyltransferase PlsX. Metab Eng 45, 59–66 (2018).

22. L. Yao, I. Cengic, J. Anfelt, E. P. Hudson, Multiple Gene Repression in Cyanobacteria Using CRISPRi. ACS Synth Biol 5, 207–212 (2016).

23. A. C. Reis et al., Simultaneous repression of multiple bacterial genes using nonrepetitive extra-long sgRNA arrays. Nat Biotechnol 37, 1294–1301 (2019).

24. J. E. McDade et al., Legionnaires’ disease: isolation of a bacterium and demonstration of its role in other respiratory disease. N Engl J Med 297, 1197–1203 (1977).

25. D. W. Fraser et al., Legionnaires’ disease: description of an epidemic of pneumonia. N Engl J Med 297, 1189–1197 (1977).

26. C. Rao et al., Active and adaptive *Legionella* CRISPR-Cas reveals a recurrent challenge to the pathogen. Cell Microbiol 18, 1319–1338 (2016).

27. K. H. Berger, R. R. Isberg, Two distinct defects in intracellular growth complemented by a single genetic locus in *Legionella pneumophila*. Mol Microbiol 7, 7–19 (1993).

28. E. Choudhary, P. Thakur, M. Pareek, N. Agarwal, Gene silencing by CRISPR interference in mycobacteria. Nat Commun 6, 6267 (2015).

29. W. Jiang, D. Bikard, D. Cox, F. Zhang, L. A. Marraffini, RNA-guided editing of bacterial genomes using CRISPR-Cas systems. Nat Biotechnol 31, 233–239 (2013).

30. D. Burstein et al., Genomic analysis of 38 *Legionella* species identifies large and diverse effector repertoires. Nat Genet 48, 167–175 (2016).

31. L. Gomez-Valero et al., More than 18,000 effectors in the Legionella genus genome provide multiple, independent combinations for replication in human cells. Proc Natl Acad Sci U S A 116, 2265–2273 (2019).

32. I. Grissa, G. Vergnaud, C. Pourcel, CRISPRFinder: a web tool to identify clustered regularly interspaced short palindromic repeats. Nucleic Acids Res 35, W52–57 (2007).

33. M. L. Luo, A. S. Mullis, R. T. Leenay, C. L. Beisel, Repurposing endogenous type I CRISPR-Cas systems for programmable gene repression. Nucleic Acids Res 43, 674–681 (2015).

34. C. Liao et al., Modular one-pot assembly of CRISPR arrays enables library generation and reveals factors influencing crRNA biogenesis. Nat Commun 10, 2948 (2019).

35. K. B. Arnvig et al., Evolutionary comparison of ribosomal operon antitermination function. J Bacteriol 190, 7251–7257 (2008).

36. P. Mitra, G. Ghosh, M. Hafeezunnisa, R. Sen, Rho Protein: Roles and Mechanisms. Annu Rev Microbiol 71, 687–709 (2017).

37. M. Torres, J. M. Balada, M. Zellars, C. Squires, C. L. Squires, In vivo effect of NusB and NusG on rRNA transcription antitermination. J Bacteriol 186, 1304–1310 (2004).

38. C. L. Squires, J. Greenblatt, J. Li, C. Condon, C. L. Squires, Ribosomal RNA antitermination in vitro: requirement for Nus factors and one or more unidentified cellular components. Proc Natl Acad Sci U S A 90, 970–974 (1993).

39. A. M. Stringer, G. Baniulyte, E. Lasek-Nesselquist, J. T. Wade, Nus Factors Prevent Premature Transcription Termination of Bacterial CRISPR Arrays. bioRxiv 10.1101/405316 (2018).

40. J. P. Vogel, R. R. Isberg, Cell biology of *Legionella pneumophila*. Curr Opin Microbiol 2, 30–34 (1999).

41. H. L. Andrews, J. P. Vogel, R. R. Isberg, Identification of linked *Legionella pneumophila* genes essential for intracellular growth and evasion of the endocytic pathway. Infect Immun 66, 950–958 (1998).

42. G. Yerushalmi, T. Zusman, G. Segal, Additive effect on intracellular growth by *Legionella pneumophila* Icm/Dot proteins containing a lipobox motif. Infect Immun 73, 7578–7587 (2005).

43. D. Chetrit, B. Hu, P. J. Christie, C. R. Roy, J. Liu, A unique cytoplasmic ATPase complex defines the *Legionella pneumophila* type IV secretion channel. Nat Microbiol 3, 678–686 (2018).

44. C. Sundstrom, K. Nilsson, Establishment and characterization of a human histiocytic lymphoma cell line (U-937). Int J Cancer 17, 565–577 (1976).

45. E. Pearlman, A. H. Jiwa, N. C. Engleberg, B. I. Eisenstein, Growth of *Legionella pneumophila* in a human macrophage-like (U937) cell line. Microb Pathog 5, 87–95 (1988).

46. D. T. Isaac, R. K. Laguna, N. Valtz, R. R. Isberg, MavN is a *Legionella pneumophila* vacuole-associated protein required for efficient iron acquisition during intracellular growth. Proc Natl Acad Sci U S A 112, E5208–5217 (2015).

47. E. T. Christenson et al., The iron-regulated vacuolar *Legionella pneumophila* MavN protein is a transition-metal transporter. Proc Natl Acad Sci U S A 116, 17775–17785 (2019).

48. A. Vigouroux, E. Oldewurtel, L. Cui, D. Bikard, S. van Teeffelen, Tuning dCas9’s ability to block transcription enables robust, noiseless knockdown of bacterial genes. Mol Syst Biol 14, e7899 (2018).

49. M. Jost et al., Titrating gene expression using libraries of systematically attenuated CRISPR guide RNAs. Nat Biotechnol 38, 355–364 (2020).

50. J. McGinn, L. A. Marraffini, Molecular mechanisms of CRISPR-Cas spacer acquisition. Nat Rev Microbiol 17, 7–12 (2019).

51. S. Ghosh, T. J. O’Connor, Beyond Paralogs: The Multiple Layers of Redundancy in Bacterial Pathogenesis. Front Cell Infect Microbiol 7, 467 (2017).

52. J. C. Feeley et al., Charcoal-yeast extract agar: primary isolation medium for *Legionella pneumophila*. J Clin Microbiol 10, 437–441 (1979).

53. M. R. Neunuebel et al., De-AMPylation of the small GTPase Rab1 by the pathogen *Legionella pneumophila*. Science 333, 453–456 (2011).

54. S. M. VanRheenen, Z. Q. Luo, T. O’Connor, R. R. Isberg, Members of a *Legionella pneumophila* family of proteins with ExoU (phospholipase A) active sites are translocated to target cells. Infect Immun 74, 3597–3606 (2006).

55. Y. H. Lin et al., RavN is a member of a previously unrecognized group of *Legionella pneumophila* E3 ubiquitin ligases. PLoS Pathog 14, e1006897 (2018).

56. G. M. Conover, I. Derre, J. P. Vogel, R. R. Isberg, The *Legionella pneumophila* LidA protein: a translocated substrate of the Dot/Icm system associated with maintenance of bacterial integrity. Mol Microbiol 48, 305–321 (2003).

57. T. D. Schmittgen, K. J. Livak, Analyzing real-time PCR data by the comparative C(T) method. Nat Protoc 3, 1101–1108 (2008).

