## Supplemental Figures for "A multiplex CRISPR interference tool for virulence gene interrogation in an intracellular pathogen"

**Supplementary Information**

**Table S1: Cloning plasmids and strains used in this study**

**Table S2: CRISPRi constructs and strains used in this study**

**Table S3: Primers used in this study**

**Figure S1: CRISPRi construct construction**

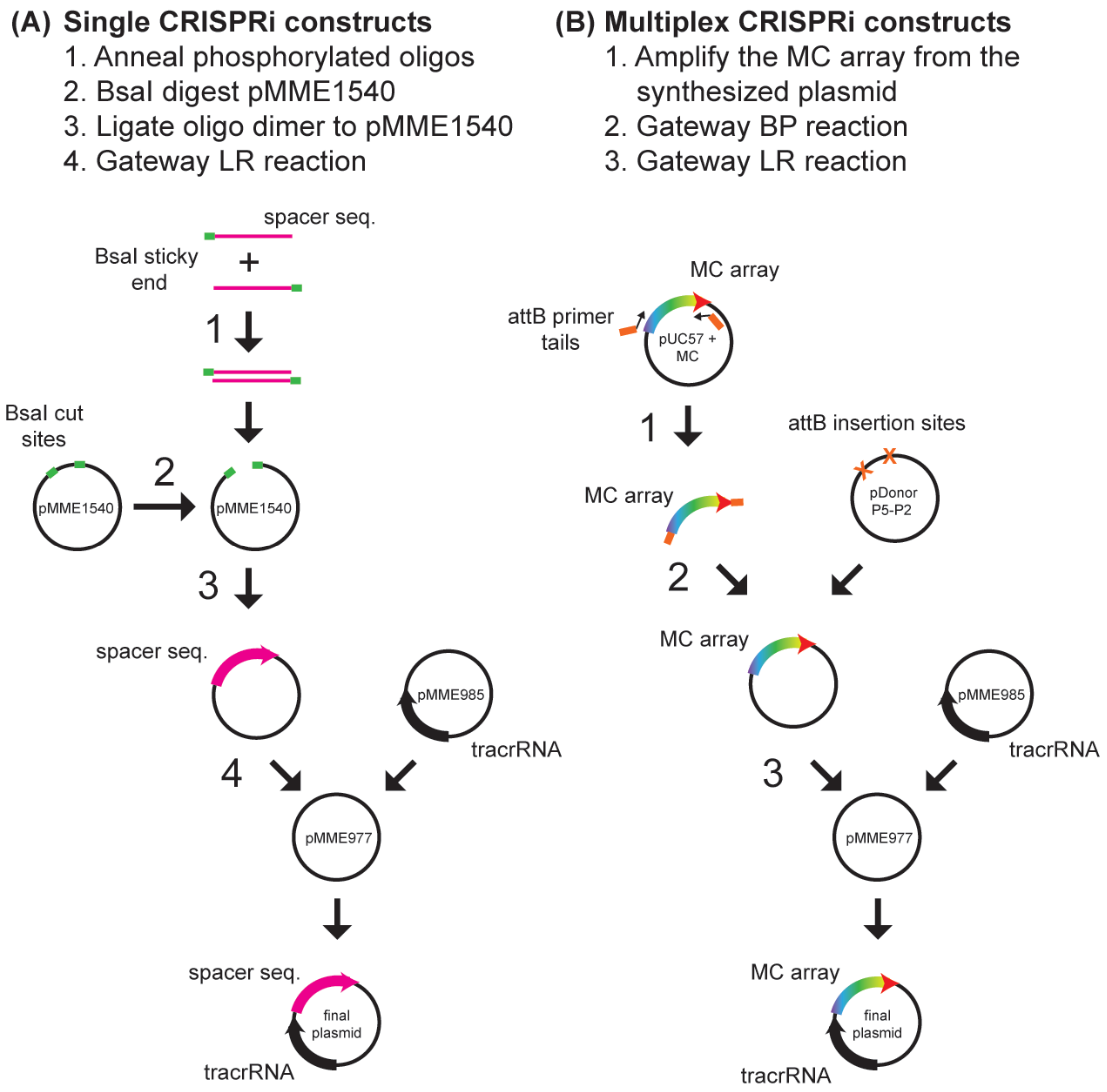


(A) Single crRNA-encoding plasmids were built using oligonucleotides and Gateway-compatible plasmids. First, a single crRNA-encoding spacer sequence was designed as complimentary oligonucleotides with BsaI-compatible overhangs, phosphorylated, and annealed as previously described for pCRISPR. Second, pMME1540 was digested with BsaI. Third, the annealed single spacer oligonucleotides were ligated into the BsaI-pre-cut pMME1540. Fourth, the inserts of this donor plasmid and pMME985, bearing the tracrRNA-encoding sequence, were transferred into pMME977 by the Invitrogen Gateway LR reaction to generate single CRISPRi plasmids. (B) Multiplex CRISPRi repeat/spacer arrays (MC arrays) were synthesized on the pUC57 vector by GenScript. First, PCR was performed using Gateway5-Ptet and Gateway2-T1term primers. Second, the multiplex CRISPRi array was introduced into pDonorP5-P2 by the Invitrogen Gateway BP reaction. Third, the inserts of this donor plasmid and pMME985, bearing the tracrRNA-encoding sequence, were transferred into pMME977 by the Invitrogen Gateway LR reaction to generate multiplex CRISPRi plasmids.

**Figure S2: Immunoblot replicates for Figure 2C**

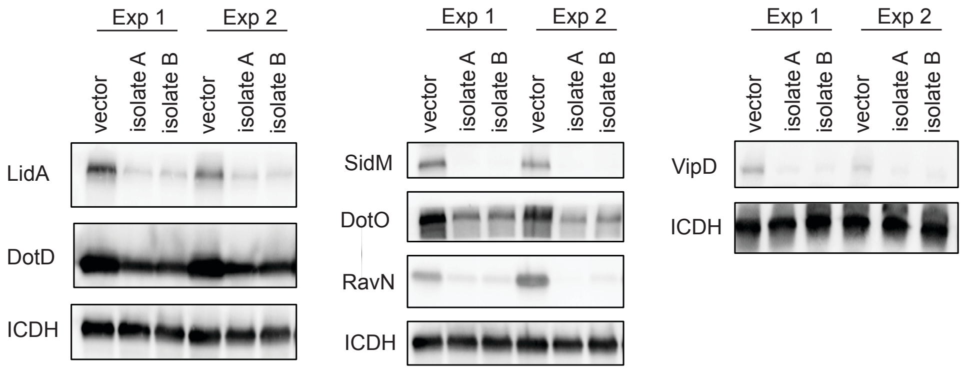


Bacteria pellets from Lp02(*dcas9*) strains bearing either the empty vector or MC-I were collected after axenic culture over the course of two separate experiments with two separate isolates (+40 ng/mL aTC). For each experiment six different proteins were probed. Multiple immunoblots were run to accommodate the range of target sizes. Correlation of target and spacer position can be found in Figure 2C. ICDH served as a loading control for each blot.

**Figure S3: Immunoblot replicates and loading controls for Figure 3C**

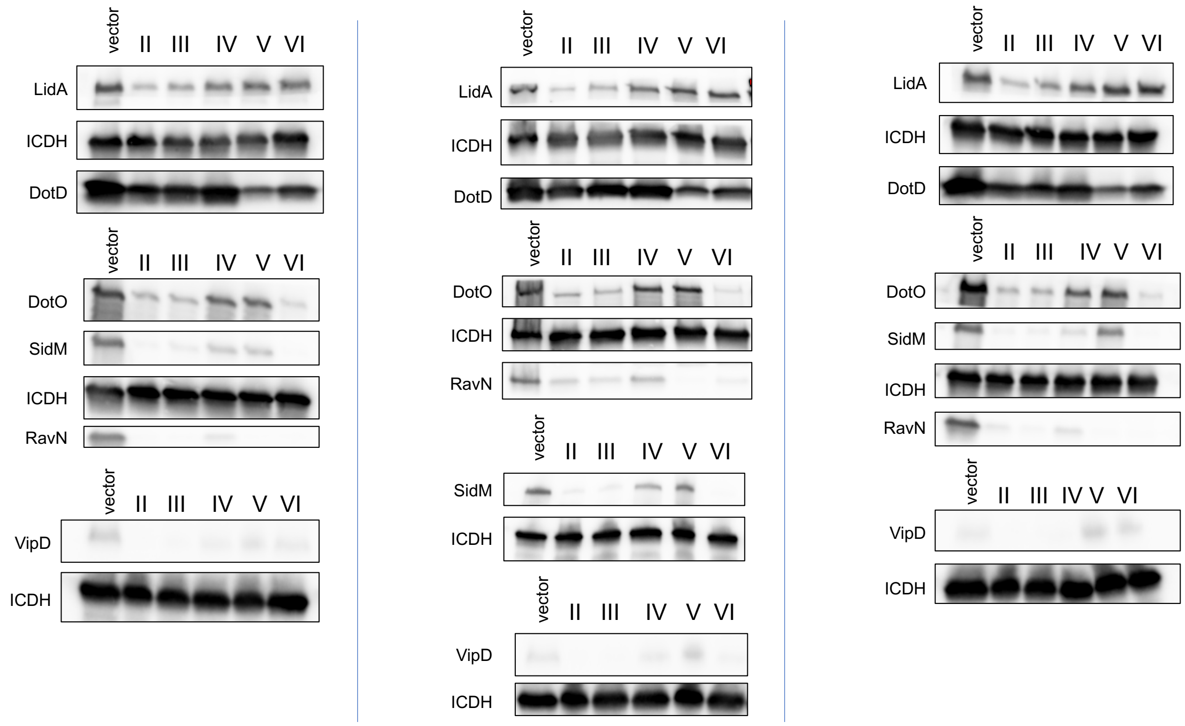


Bacteria pellets from Lp02(*dcas9*) bearing either the empty vector or a $P_{tet}$-MC construct were collected after axenic culture over the course of three separate experiments (+40 ng/mL aTC). For each experiment six different proteins were probed. Multiple immunoblots were run to accommodate the range of target sizes. Roman numerals correspond to the $P_{tet}$-MC construct number. Correlation of construct and spacer position can be found in Figure 3A. ICDH served as a loading control for each blot.

**Figure S4: Upstream region of *boxA* constructs**

**Promoter-leader-1st repeat**

```
tccctatcagtgatagagattgacatccctatcagtgatagagatactgagcacaataaatg
cagtaatacaggggcttttcaagactgaagtctagctgagacaaatagtgcgattacgaaat
tttttagacaaaaatagtctacgaggttttagagctatgctgttttgaatggtcccaaaac
```

***boxA*(-58)**

```
tccctatcagtgatagagattgacatccctatcagtgatagagatactgagcacaataaatg
cagtaatacaggggcttttcaagactgaaATTTAGTTCTTTAAAAATTTGcgattacgaaat
tttttagacaaaaatagtctacgaggttttagagctatgctgttttgaatggtcccaaaac
```

***boxA*(-90)**

```
tccctatcagtgatagagattgacatccctatcagtgatagagatactgagcacaataaATT
TAGTTCTTTAAAAATTTTtcaagactgaagtctagctgagacaaatagtgcgattacgaaat
tttttagacaaaaatagtctacgaggttttagagctatgctgttttgaatggtcccaaaac
```

The promoter (blue), leader (black), and first repeat (yellow) sequence are shown for the original *boxA*-less $P_{tet}$-MC constructs. Below, the sequence was mutated to include *boxA* (pink underline with five flanking base pairs) from the *L. pneumophila sp. Lens* CRISPR array at either -58 base pairs or -90 base pairs upstream of the first repeat.

**Figure S5: Immunoblot replicates and loading controls for Figure 4B**

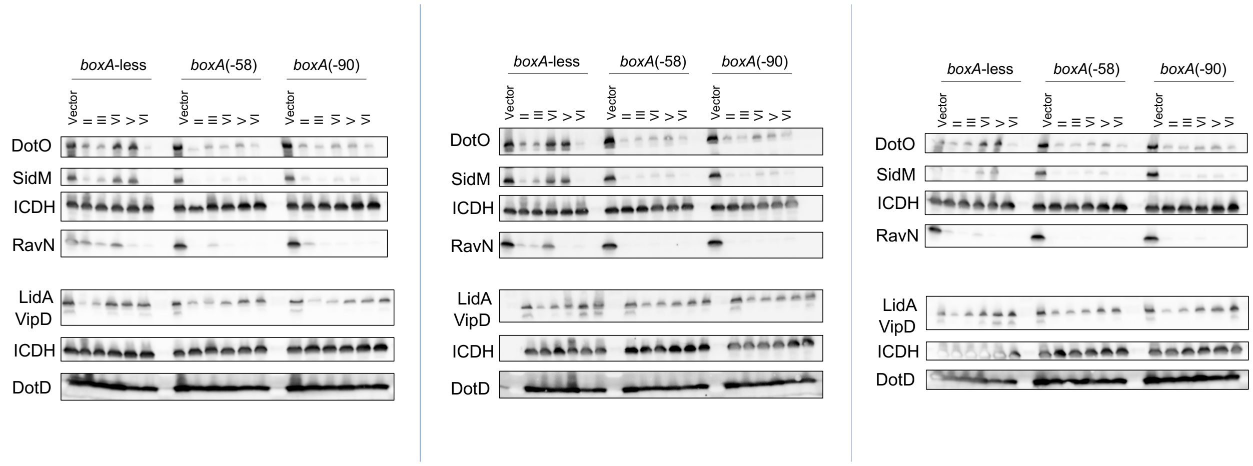


Bacteria pellets from Lp02(*dcas9*) bearing either the empty vector or a *boxA*-less, *boxA*(-58), or *boxA*(-90) $P_{tet}$-MC construct were collected after axenic culture over the course of three separate experiments (+40 ng/mL aTC). For each experiment six different proteins were probed. Multiple immunoblots were run to accommodate the range of target sizes. Roman numerals correspond to the MC construct number. Correlation of construct and spacer position can be found in Figure 3A. ICDH served as a loading control for each blot.

**Figure S6: Immunoblot replicates and loading controls for Figure 6C**

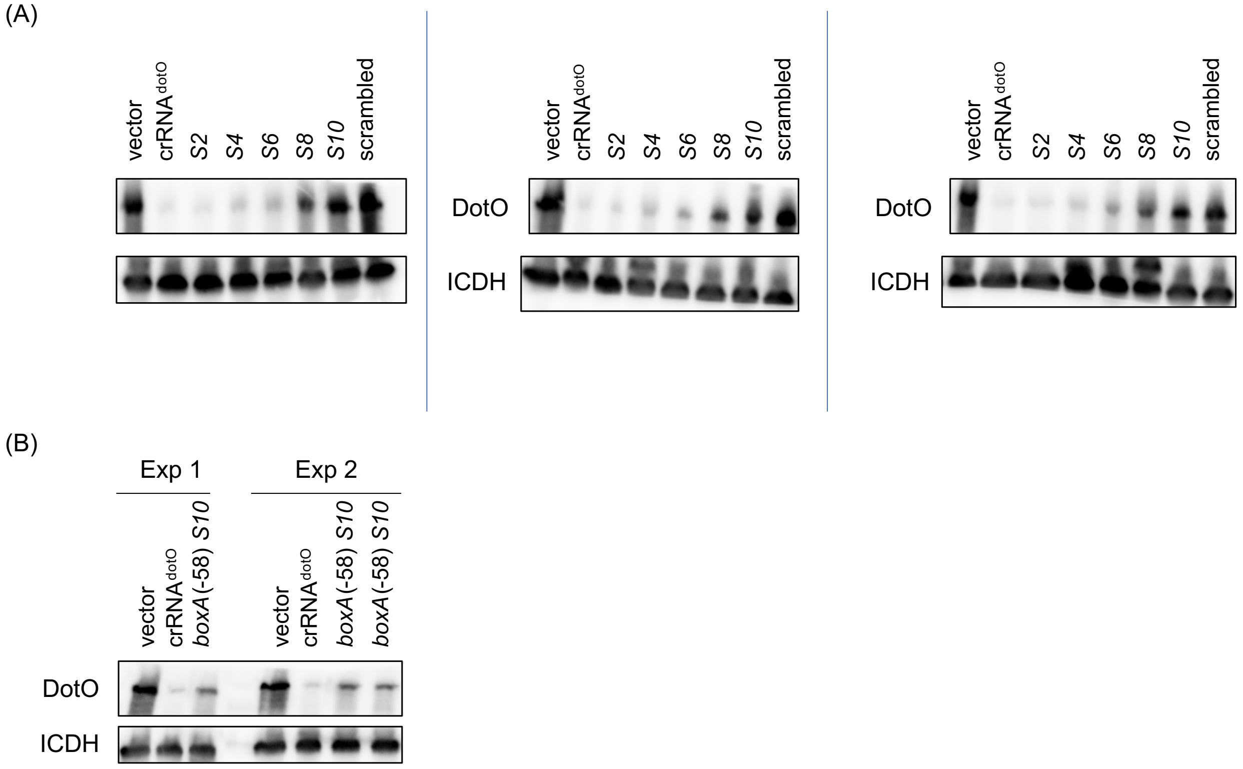


Immunoblot analyses were performed on bacteria pellets collected from axenic cultures of (A) *boxA*-less and (B) *boxA*(-58) $P_{tet}$-MC-dotO constructs and the empty vector Lp02(*dcas9*) strains (+40 ng/mL aTC). Samples were organized according to *dotO* spacer position. Correlation of spacer position and DotO construct can be found in Figure 6A. ICDH serves as a loading control.

### Figure S7: Optimized 10-plex CRISPRi templates

**(A) *boxA-less* $P_{tet}$-MC**

**$P_{tet}$ promoter-leader-[repeat-spacer(“n”s)]-extra-T1 terminator**

```
tccctatcagtgatagagattgacatccctatcagtgatagagatactgagcac
aataaatgcagtaatacaggggcttttcaagactgaagtctagctgagacaaat
agtgcgattacgaaatttttttagacaaaaatagtctacgag
gttttagagctatgctgttttgaatggtcccaaaac
nnnnnnnnnnnnnnnnnnnnnnnnnnnnnn
gttttagagctatgctgttttgaatggtcccaaaac
nnnnnnnnnnnnnnnnnnnnnnnnnnnnnn
gttttagagctatgctgttttgaatggtcccaaaac
nnnnnnnnnnnnnnnnnnnnnnnnnnnnnn
gttttagagctatgctgttttgaatggtcccaaaac
nnnnnnnnnnnnnnnnnnnnnnnnnnnnnn
gttttagagctatgctgttttgaatggtcccaaaac
nnnnnnnnnnnnnnnnnnnnnnnnnnnnnn
gttttagagctatgctgttttgaatggtcccaaaac
nnnnnnnnnnnnnnnnnnnnnnnnnnnnnn
gttttagagctatgctgttttgaatggtcccaaaac
nnnnnnnnnnnnnnnnnnnnnnnnnnnnnn
gttttagagctatgctgttttgaatggtcccaaaac
nnnnnnnnnnnnnnnnnnnnnnnnnnnnnn
gttttagagctatgctgttttgaatggtcccaaaac
nnnnnnnnnnnnnnnnnnnnnnnnnnnnnn
gttttagagctatgctgttttgaatggtcccaaaac
nnnnnnnnnnnnnnnnnnnnnnnnnnnnnn
gttttagagctatgctgttttgaatggtcccaaaac
ttcagcacactgagacttgttgagttgaattc
gggaactgccaggcatcaaataaaacgaaaggctcagtcgaaagactgggccttt
cgttttatctgttgtttgtcggtgaacgctctcctg
```

**(B) *boxA*(-58) $P_{tet}$-MC**

**$P_{tet}$ promoter-leader with *boxA(-58)*-[repeat-spacer(“n”s)]-extra-T1 terminator**

```
tccctatcagtgatagagattgacatccctatcagtgatagagatactgagcac
aataaatgcagtaatacaggggcttttcaagactgaaATTTAGTTCTTTAAAAA
TTTGcgattacgaaatttttttagacaaaaatagtctacgag
gttttagagctatgctgttttgaatggtcccaaaac
nnnnnnnnnnnnnnnnnnnnnnnnnnnnnn
gttttagagctatgctgttttgaatggtcccaaaac
nnnnnnnnnnnnnnnnnnnnnnnnnnnnnn
gttttagagctatgctgttttgaatggtcccaaaac
nnnnnnnnnnnnnnnnnnnnnnnnnnnnnn
gttttagagctatgctgttttgaatggtcccaaaac
nnnnnnnnnnnnnnnnnnnnnnnnnnnnnn
gttttagagctatgctgttttgaatggtcccaaaac
nnnnnnnnnnnnnnnnnnnnnnnnnnnnnn
gttttagagctatgctgttttgaatggtcccaaaac
nnnnnnnnnnnnnnnnnnnnnnnnnnnnnn
gttttagagctatgctgttttgaatggtcccaaaac
nnnnnnnnnnnnnnnnnnnnnnnnnnnnnn
gttttagagctatgctgttttgaatggtcccaaaac
nnnnnnnnnnnnnnnnnnnnnnnnnnnnnn
gttttagagctatgctgttttgaatggtcccaaaac
nnnnnnnnnnnnnnnnnnnnnnnnnnnnnn
gttttagagctatgctgttttgaatggtcccaaaac
nnnnnnnnnnnnnnnnnnnnnnnnnnnnnn
gttttagagctatgctgttttgaatggtcccaaaac
ttcagcacactgagacttgttgagttgaattc
gggaactgccaggcatcaaataaaacgaaaggctcagtcgaaagactgggccttt
cgttttatctgttgtttgtcggtgaacgctctcctg
```

Nucleotide sequence for multiplex CRISPRi platforms (A) *boxA*-less Ptet-MC or (B) *boxA*(-58) $P_{tet}$-MC. Repeat sequences are 36 base pairs long and spacer sequences are 30 base pairs long (represented by 30 “n”s). Spacer sequences used in this study can be found in Table S2.
