## Supplemental Table 1 for "A multiplex CRISPR interference tool for virulence gene interrogation in an intracellular pathogen"

### Table S1: Cloning plasmids and strains used in this study

| *L. pneumophila* Strains | Description | Selection | Reference |
|---|---|---|---|
| Lp02 | Philadelphia 1, *thyA rpsL, hsdR* | thymidine auxotroph | Berger KH, Isberg RR (1993) Two distinct defects in intracellular growth complemented by a single genetic locus in *Legionella pneumophila*. *Mol Microbiol* 7:7-19. |
| Lp03 | *thyA rpsL hsdR dotA03* | thymidine auxotroph | Berger KH, Isberg RR (1993) Two distinct defects in intracellular growth complemented by a single genetic locus in *Legionella pneumophila*. *Mol Microbiol* 7:7-19. |
| MML109 (Lp02(*dcas9*)) | Lp02 with *dcas9* on the chromosome at *thyA* | thymidine auxotroph | This study |

| *E. coli* Strains | Description | Selection | Reference |
|---|---|---|---|
| GC5 | GC5 Competent Cells | | Genesee Scientific (Catalog #42-650) |
| PIR2+ | One Shot™ PIR2 Chemically Competent *E. coli* | | Invitrogen (Catalog #C111110) |

| Plasmids | Description | Selection | Reference |
|---|---|---|---|
| pdCas9-bacteria | aTc-inducible expression of a catalytically inactive bacterial Cas9 (*S. pyogenes*) | Cm$^R$ | Addgene plasmid #44249 |
| pCRISPR | a crRNA expression plasmid for targeting a specific sequence | Kan$^R$ | Addgene plasmid #42875 |
| pCas9 | for bacterial expression of Cas9 nuclease, tracrRNA and crRNA guide | Cm$^R$ | Addgene plasmid #42876 |
| pNTPS138_Cm | *sacB* bacterial exchange vector | Cm$^R$ | Addgene plasmid #41891 |
| pJB908 | *Legionella* expression vector; a derivative of pKB5 (AmpR *td* Δi) | Amp$^R$ | Hammer BK, Swanson MS (1999) Co-ordination of *Legionella pneumophila* virulence with entry into stationary phase by ppGpp. *Mol Microbiol* 33(4):721-31. |
| pDonorP4r-P3r | a Gateway entry vector | Kan$^R$ | Invitrogen MultiSite Gateway ProPlus (Catalog #45-2100) |
| pDonorP1-P4 | a Gateway entry vector | Kan$^R$ | Invitrogen MultiSite Gateway ProPlus (Catalog #45-2100) |
| pDonorP3-P2 | a Gateway entry vector | Kan$^R$ | Invitrogen MultiSite Gateway ProPlus (Catalog #45-2100) |
| pDonorP5-P2 | a Gateway entry vector | Kan$^R$ | Invitrogen MultiSite Gateway ProPlus (Catalog #45-2100) |
| pDonorP1-P5 | a Gateway entry vector | Kan$^R$ | Invitrogen MultiSite Gateway ProPlus (Catalog #45-2100) |
| pDEST17 | a Gateway destination vector | Amp$^R$ | Invitrogen (Catalog #11803012) |
| pMME1080 | dCas9 amplified from pdCas9-bacteria in pDonorP4r-P3r | Kan$^R$ | This study |
| pMME1084 | homologous recombinant region of N-terminus *thyA* in pDonorP1-P4 | Kan$^R$ | This study |
| pMME1094 | homologous recombinant region of C-terminus *thyA* in pDonorP3-P2 | Kan$^R$ | This study |
| pMME1020 | pNTPS138_Cm-DEST (Gataway compatible pNTPS138_Cm) | Cm$^R$ | This study |
| pMME1115 | *thyA*::tetR-*dcas9* cassette::C-terminus *thyA* in pNTPS138 Cm | Cm$^R$ | This study |
| pMME1540 | CRISPR-BsaI fragment amplified from pCRISPR in pDonorP5-P2 | Kan$^R$ | This study |
| pMME1748 | CRISPR-BsaI fragment with Ptet in pDonorP5-P2 | Kan$^R$ | This study |
| pMME985 | tracrDNA amplified from pCas9 in pDonorP1-P5 | Kan$^R$ | This study |
| pMME977 | *L. pneumophilila* compatible destination vector(pJB908D) | Amp$^R$ | Lin Y, Doms AG, Cheng E, Kim B, Evans TR, Machner MP (2015) Host cell-catalyzed S-palmitoylation mediates Golgi targeting of the Legionella ubiquitin ligase GobX. J Biol Chem 290 (42): 25766-81 |
| pMME1568 | LidA single crRNA in pMME1540 | Kan$^R$ | This study |
| pMME998 | SidM single crRNA - A in pMME1540 | Kan$^R$ | This study |
| pMME1059 | SidM single crRNA - B in pMME1540 | Kan$^R$ | This study |
| pMME1062 | SidM single crRNA - C in pMME1540 | Kan$^R$ | This study |
| pMME1063 | SidM single crRNA - D in pMME1540 | Kan$^R$ | This study |
| pMME1000 | VipD single crRNA - A in pMME1540 | Kan$^R$ | This study |
| pMME1073 | VipD single crRNA - B in pMME1540 | Kan$^R$ | This study |
| pMME1067 | VipD single crRNA - C in pMME1540 | Kan$^R$ | This study |
| pMME1069 | VipD single crRNA - D in pMME1540 | Kan$^R$ | This study |
| pMME1717 | DotO single crRNA in pMME1540 | Kan$^R$ | This study |
| pMME1170 | Mlong-pUC57 (21 crRNA encoding sequences) | Amp$^R$ | Synthesized by GenScript for this study |
| pMME1571 | Pnative-MC-3/4 in pMME1540 | Kan$^R$ | This study |
| pMME1575 | Pnative-MC-7/8 in pMME1540 | Kan$^R$ | This study |
| pMME1577 | Pnative-MC-9/10 in pMME1540 | Kan$^R$ | This study |
| pMME1583 | Ptet-MC-3/4 in pMME1540 | Kan$^R$ | This study |
| pMME1585 | Ptet-MC-7/8 in pMME1540 | Kan$^R$ | This study |
| pMME1586 | Ptet-MC-9/10 in pMME1540 | Kan$^R$ | This study |
| pMME2355 | MC-I in pUC57 | Amp$^R$ | Synthesized by GenScript for this study |
| pMME2356 | MC-II in pUC57 | Amp$^R$ | Synthesized by GenScript for this study |
| pMME2357 | MC-III in pUC57 | Amp$^R$ | Synthesized by GenScript for this study |
| pMME2358 | MC-IV in pUC57 | Amp$^R$ | Synthesized by GenScript for this study |
| pMME2359 | MC-V in pUC57 | Amp$^R$ | Synthesized by GenScript for this study |
| pMME2360 | MC-VI in pUC57 | Amp$^R$ | Synthesized by GenScript for this study |
| pMME2367 | MC-I in pDonorP5-P2 (boxA-less) | Kan$^R$ | This study |
| pMME2368 | MC-II in pDonorP5-P2 (boxA-less) | Kan$^R$ | This study |
| pMME2369 | MC-III in pDonorP5-P2 (boxA-less) | Kan$^R$ | This study |
| pMME2370 | MC-IV in pDonorP5-P2 (boxA-less) | Kan$^R$ | This study |
| pMME2371 | MC-V in pDonorP5-P2 (boxA-less) | Kan$^R$ | This study |
| pMME2372 | MC-VI in pDonorP5-P2 (boxA-less) | Kan$^R$ | This study |
| pMME2373 | MC-II in pDonorP5-P2 (boxA(-58)) | Kan$^R$ | This study |
| pMME2374 | MC-III in pDonorP5-P2 (boxA(-58)) | Kan$^R$ | This study |
| pMME2375 | MC-IV in pDonorP5-P2 (boxA(-58)) | Kan$^R$ | This study |
| pMME2376 | MC-V in pDonorP5-P2 (boxA(-58)) | Kan$^R$ | This study |
| pMME2377 | MC-VI in pDonorP5-P2 (boxA(-58)) | Kan$^R$ | This study |
| pMME2383 | MC-II in pDonorP5-P2 (boxA(-90)) | Kan$^R$ | This study |
| pMME2384 | MC-III in pDonorP5-P2 (boxA(-90)) | Kan$^R$ | This study |
| pMME2385 | MC-IV in pDonorP5-P2 (boxA(-90)) | Kan$^R$ | This study |
| pMME2386 | MC-V in pDonorP5-P2 (boxA(-90)) | Kan$^R$ | This study |
| pMME2387 | MC-VI in pDonorP5-P2 (boxA(-90)) | Kan$^R$ | This study |
| pMME2393 | DotO-II in pUC57 | Amp$^R$ | Synthesized by GenScript for this study |
| pMME2394 | DotO-III in pUC57 | Amp$^R$ | Synthesized by GenScript for this study |
| pMME2395 | DotO-IV in pUC57 | Amp$^R$ | Synthesized by GenScript for this study |
| pMME2396 | DotO-V in pUC57 | Amp$^R$ | Synthesized by GenScript for this study |
| pMME2579 | DotO-VI in pUC57 | Amp$^R$ | Synthesized by GenScript for this study |
| pMME2580 | DotO-VII in pUC57 | Amp$^R$ | Synthesized by GenScript for this study |
| pMME2581 | DotO-II in pDonorP5-P2 (boxA-less) | Kan$^R$ | This study |

| | | | |
|---|---|---|---|
| pMME2582 | DotO-III in pDonorP5-P2 (boxA-less) | $Kan^R$ | This study |
| pMME2583 | DotO-IV in pDonorP5-P2 (boxA-less) | $Kan^R$ | This study |
| pMME2584 | DotO-V in pDonorP5-P2 (boxA-less) | $Kan^R$ | This study |
| pMME2585 | DotO-VI in pDonorP5-P2 (boxA-less) | $Kan^R$ | This study |
| pMME2586 | DotO-VII in pDonorP5-P2 (boxA-less) | $Kan^R$ | This study |
| pMME2590 | DotO-V in pDonorP5-P2 (boxA(-58)) | $Kan^R$ | This study |
