## Supplemental Table 2 for "A multiplex CRISPR interference tool for virulence gene interrogation in an intracellular pathogen"

### Table S2: CRISPRi constructs and strains used in this study

| Name | Spacer position | Targeted gene | crRNA sequence 5' to 3' | CRISPR construct (AmpR) | *Legionella* strains bearing CRISPR construct | |
|---|---|---|---|---|---|---|
| | | | | | Lp02(-*dcas9*) | Lp02(*dcas9*) = MML109 |
| LidA single crRNA | 1 | *lpg0940* | ctaaaacaggttccttttccttggttttta | pMME1581 | MML283 | MML284 |
| SidM single crRNA - A | 1 | *lpg2464* | acttatgaatcctcgagtacatgagcataa | pMME1516 | MML154 | MML155 |
| SidM single crRNA - B | 1 | *lpg2464* | aattaaaatgagtgttaatgaagagcaatt | pMME1518 | MML156 | MML157 |
| SidM single crRNA - C | 1 | *lpg2464* | aaatttttttctgagcggttggggatagtaa | pMME1130 | MML126 | MML127 |
| SidM single crRNA - D | 1 | *lpg2464* | atctcgtaactcttctcctaattcatagcg | pMME1519 | MML158 | MML159 |
| VipD single crRNA - A | 1 | *lpg2831* | tttaaacaaacaattatgtgagcaattgat | pMME1517 | MML183 | MML184 |
| VipD single crRNA - B | 1 | *lpg2831* | aaaggaatttactggaaatatcaaagactg | pMME1522 | MML189 | MML190 |
| VipD single crRNA - C | 1 | *lpg2831* | gatatttccagtaaattcctttttaattta | pMME1521 | MML185 | MML186 |
| VipD single crRNA - D | 1 | *lpg2831* | gccataattcatctgttgttctggaggaat | pMME1538 | MML187 | MML188 |
| DotO single crRNA | 1 | *lpg0456* | AATCTATATAAGACTCTGTAGTTTGCTTCA | pMME1730 | | MML266 |
| DotD single crRNA | 1 | *lpg2674* | GGTCGCATCATCACTTGGATTATTGATGGG | pMME1725 | | MML263 |
| MavN single crRNA | 1 | *lpg2815* | CAAGTAATAAGGGATTTTTTGGGGCTTAAT | pMME1747 | MML279 | MML280 |
| MC-3/4 | 1 | *lpg2584 (sidF)* | GTTCCAGATCACTTAATAAATGCGTGACAT | Pnative: pMME1562 | Pnative: MML199 | Pnative: MML200 |
| | 2 | | TAAGTATTTACTTTTATGATCAATATCAAA | Ptet: pMME1596 | Ptet: MML218 | Ptet: MML219 |
| | 3 | *lpg0940 (lidA)* | CTAAAACAGGTTCCTTTTCCTTGGTTTTTA | | | |
| | 4 | *lpg0940 (lidA)* | AAATACCACAAAAACAATGCGGCCATTAAT | | | |
| MC-7/8 | 1 | *lpg2465(sidD)* | AGATTTCACTATTACCCTTATCTAAATCAT | Pnative: pMME1564 | Pnative: MML209 | Pnative: MML210 |
| | 2 | | TAATTCCGTGAATAGTTCACGAGGGTCACC | Ptet: pMME1600 | Ptet: MML222 | Ptet: MML223 |
| | 3 | | TTCATCAACATATAGATAATCACAACGTTC | | | |
| | 4 | | AAATTTTCTTCTTAAAACCATTATCACTTC | | | |
| | 5 | *lpg2584 (sidF)* | GTTCCAGATCACTTAATAAATGCGTGACAT | | | |
| | 6 | | TAAGTATTTACTTTTATGATCAATATCAAA | | | |
| | 7 | *lpg0940 (lidA)* | CTAAAACAGGTTCCTTTTCCTTGGTTTTTA | | | |
| | 8 | *lpg0940 (lidA)* | AAATACCACAAAAACAATGCGGCCATTAAT | | | |
| MC-9/10 | 1 | *lpg1978 (setA)* | TTTGTCAATTTGAGCCTCTTGACCAGCCTG | Pnative: pMME1567 | Pnative: MML211 | Pnative: MML212 |
| | 2 | | TAAATGCTTTTTCGTAATACCCCTTGTTAT | Ptet: pMME1602 | Ptet: MML224 | Ptet: MML225 |
| | 3 | *lpg2465(sidD)* | AGATTTCACTATTACCCTTATCTAAATCAT | | | |
| | 4 | | TAATTCCGTGAATAGTTCACGAGGGTCACC | | | |
| | 5 | | TTCATCAACATATAGATAATCACAACGTTC | | | |
| | 6 | | AAATTTTCTTCTTAAAACCATTATCACTTC | | | |
| | 7 | *lpg2584 (sidF)* | GTTCCAGATCACTTAATAAATGCGTGACAT | | | |
| | 8 | | TAAGTATTTACTTTTATGATCAATATCAAA | | | |
| | 9 | *lpg0940 (lidA)* | CTAAAACAGGTTCCTTTTCCTTGGTTTTTA | | | |
| | 10 | *lpg0940 (lidA)* | AAATACCACAAAAACAATGCGGCCATTAAT | | | |
| MC empty vector | | *none, CRISPR array leader sequence* | AATAAATGCAGTAATACAGGGGCTTTTCAAGACTGAAGTCTAGCTGAGACAAAT AGTGCGATTACGAAATTTTTTAGACAAAAATAGTCTACGAG | pMME1936 | | MML414 |
| MC-I | 1 | *lpg0940 (lidA)* | ctaaaacaggttccttttccttggttttta | *boxA*-less: pMME2361 | | *boxA*-less: MML900 |
| | 2 | | CACCTCCTTGTAAAGCAAGATTAATCATTT | | | |
| | 3 | *lpg2464 (sidM)* | aaatttttttctgagcggttggggatagtaa | | | |
| | 4 | *lpg0208* | CTCTCGCATGCCAAATAGCTGCTCCCAAGA | | | |
| | 5 | *lpg2674 (dotD)* | GGTCGCATCATCACTTGGATTATTGATGGG | | | |
| | 6 | *lpg2831 (vipD)* | gatatttccagtaaattcctttttaattta | | | |
| | 7 | *lpg2793* | TTAAATTTAAGGGGTTAATTGGTGGAGTCA | | | |
| | 8 | *lpg0456 (dotO)* | AATCTATATAAGACTCTGTAGTTTGCTTCA | | | |
| | 9 | | CATTCATAGCACCAGCACTCGTTGCAGAAA | | | |
| | 10 | *lpg1111 (ravN)* | ATCTTCTTGCATAATAGGATCAAAATAAGT | | | |
| MC-II | 1 | *lpg0940 (lidA)* | ctaaaacaggttccttttccttggttttta | *boxA*-less: pMME2362 | | *boxA*-less: MML901 |
| | 2 | *lpg2831 (vipD)* | gatatttccagtaaattcctttttaattta | *boxA* (-58): pMME2378 | | *boxA* (-58): MML906 |
| | 3 | *lpg2464 (sidM)* | aaatttttttctgagcggttggggatagtaa | *boxA* (-90): pMME2388 | | *boxA* (-90): MML911 |
| | 4 | *lpg0456 (dotO)* | AATCTATATAAGACTCTGTAGTTTGCTTCA | | | |
| | 5 | *lpg2674 (dotD)* | GGTCGCATCATCACTTGGATTATTGATGGG | | | |
| | 6 | *lpg1111 (ravN)* | ATCTTCTTGCATAATAGGATCAAAATAAGT | | | |
| | 7 | *lpg2793* | TTAAATTTAAGGGGTTAATTGGTGGAGTCA | | | |
| | 8 | | CACCTCCTTGTAAAGCAAGATTAATCATTT | | | |
| | 9 | | CATTCATAGCACCAGCACTCGTTGCAGAAA | | | |
| | 10 | *lpg0208* | CTCTCGCATGCCAAATAGCTGCTCCCAAGA | | | |
| MC-III | 1 | | CATTCATAGCACCAGCACTCGTTGCAGAAA | *boxA*-less: pMME2363 | | *boxA*-less: MML902 |
| | 2 | *lpg0208* | CTCTCGCATGCCAAATAGCTGCTCCCAAGA | *boxA* (-58): pMME2379 | | *boxA* (-58): MML907 |
| | 3 | *lpg0940 (lidA)* | ctaaaacaggttccttttccttggttttta | *boxA* (-90): pMME2389 | | *boxA* (-90): MML912 |
| | 4 | *lpg2831 (vipD)* | gatatttccagtaaattcctttttaattta | | | |
| | 5 | *lpg2464 (sidM)* | aaatttttttctgagcggttggggatagtaa | | | |
| | 6 | *lpg0456 (dotO)* | AATCTATATAAGACTCTGTAGTTTGCTTCA | | | |
| | 7 | *lpg2674 (dotD)* | GGTCGCATCATCACTTGGATTATTGATGGG | | | |
| | 8 | *lpg1111 (ravN)* | ATCTTCTTGCATAATAGGATCAAAATAAGT | | | |
| | 9 | *lpg2793* | TTAAATTTAAGGGGTTAATTGGTGGAGTCA | | | |
| | 10 | | CACCTCCTTGTAAAGCAAGATTAATCATTT | | | |
| MC-IV | 1 | *lpg2793* | TTAAATTTAAGGGGTTAATTGGTGGAGTCA | *boxA*-less: pMME2364 | | *boxA*-less: MML903 |

| | | | | | |
|---|---|---|---|---|---|
| | 2 | | CACCTCCTTGTAAAGCAAGATTAATCATTT | *boxA* (-58): pMME2380 | *boxA* (-58): MML908 |
| | 3 | | CATTCATAGCACCAGCACTCGTTGCAGAAA | *boxA* (-90): pMME2390 | *boxA* (-90): MML913 |
| | 4 | *lpg0208* | CTCTCGCATGCCAAATAGCTGCTCCCAAGA | | |
| | 5 | *lpg0940 (lidA)* | ctaaaacaggttccttttccttggttttta | | |
| | 6 | *lpg2831 (vipD)* | gatatttccagtaaattcctttttaattta | | |
| | 7 | *lpg2464 (sidM)* | aaattttttctgagcggttggggatagtaa | | |
| | 8 | *lpg0456 (dotO)* | AATCTATATAAGACTCTGTAGTTTGCTTCA | | |
| | 9 | *lpg2674 (dotD)* | GGTCGCATCATCACTTGGATTATTGATGGG | | |
| | 10 | *lpg1111 (ravN)* | ATCTTCTTGCATAATAGGATCAAAATAAGT | | |
| MC-V | 1 | *lpg2674 (dotD)* | GGTCGCATCATCACTTGGATTATTGATGGG | *boxA* -less: pMME2365 | *boxA* -less: MML904 |
| | 2 | *lpg1111 (ravN)* | ATCTTCTTGCATAATAGGATCAAAATAAGT | *boxA* (-58): pMME2381 | *boxA* (-58): MML909 |
| | 3 | *lpg2793* | TTAAATTTAAGGGGTTAATTGGTGGAGTCA | *boxA* (-90): pMME2391 | *boxA* (-90): MML914 |
| | 4 | | AACGCTGTCCTTATAATAGAGCATTTTATC | | |
| | 5 | | CACCTCCTTGTAAAGCAAGATTAATCATTT | | |
| | 6 | *lpg0208* | CTCTCGCATGCCAAATAGCTGCTCCCAAGA | | |
| | 7 | *lpg0940 (lidA)* | ctaaaacaggttccttttccttggttttta | | |
| | 8 | *lpg2831 (vipD)* | gatatttccagtaaattcctttttaattta | | |
| | 9 | *lpg2464 (sidM)* | aaattttttctgagcggttggggatagtaa | | |
| | 10 | *lpg0456 (dotO)* | AATCTATATAAGACTCTGTAGTTTGCTTCA | | |
| MC-VI | 1 | *lpg2464 (sidM)* | aaattttttctgagcggttggggatagtaa | *boxA* -less: pMME2366 | *boxA* -less: MML905 |
| | 2 | *lpg0456 (dotO)* | AATCTATATAAGACTCTGTAGTTTGCTTCA | *boxA* (-58): pMME2382 | *boxA* (-58): MML910 |
| | 3 | *lpg2674 (dotD)* | GGTCGCATCATCACTTGGATTATTGATGGG | *boxA* (-90): pMME2392 | *boxA* (-90): MML915 |
| | 4 | *lpg1111 (ravN)* | ATCTTCTTGCATAATAGGATCAAAATAAGT | | |
| | 5 | *lpg2793* | TTAAATTTAAGGGGTTAATTGGTGGAGTCA | | |
| | 6 | | AACGCTGTCCTTATAATAGAGCATTTTATC | | |
| | 7 | | CATTCATAGCACCAGCACTCGTTGCAGAAA | | |
| | 8 | *lpg0208* | CTCTCGCATGCCAAATAGCTGCTCCCAAGA | | |
| | 9 | *lpg0940 (lidA)* | ctaaaacaggttccttttccttggttttta | | |
| | 10 | *lpg2831 (vipD)* | gatatttccagtaaattcctttttaattta | | |
| MC-II-dotO | 1 | | ctctttcgtttattatcaagtacttagtcg | *boxA* -less: pMME2592 | *boxA* -less: MML916 |
| | 2 | | Atcttaactttaataatatttattctgtcg | | |
| | 3 | | Tagttgaagtttctgggaaatatgcgatgt | | |
| | 4 | *lpg0456 (dotO)* | AATCTATATAAGACTCTGTAGTTTGCTTCA | | |
| | 5 | | GTATAACCGCGTACTTGTAGCGTGAGTGTT | | |
| | 6 | | ATAACCGGATATTAATTAGGTTTAAACTAC | | |
| | 7 | | GTAGAAGTGGGTTTATTCGTAAAGATGTTA | | |
| | 8 | | AGATATCTGCCATATTATACAAACGTTTCC | | |
| | 9 | | CGAGTGACCGGTCTCACACAACTACAATTA | | |
| | 10 | | CACGCCGCATATCGGAATCCACTATCTGCA | | |
| MC-III-dotO | 1 | | CGAGTGACCGGTCTCACACAACTACAATTA | *boxA* -less: pMME2593 | *boxA* -less: MML917 |
| | 2 | | CACGCCGCATATCGGAATCCACTATCTGCA | | |
| | 3 | | ctctttcgtttattatcaagtacttagtcg | | |
| | 4 | | Atcttaactttaataatatttattctgtcg | | |
| | 5 | | Tagttgaagtttctgggaaatatgcgatgt | | |
| | 6 | *lpg0456 (dotO)* | AATCTATATAAGACTCTGTAGTTTGCTTCA | | |
| | 7 | | GTATAACCGCGTACTTGTAGCGTGAGTGTT | | |
| | 8 | | ATAACCGGATATTAATTAGGTTTAAACTAC | | |
| | 9 | | GTAGAAGTGGGTTTATTCGTAAAGATGTTA | | |
| | 10 | | AGATATCTGCCATATTATACAAACGTTTCC | | |
| MC-IV-dotO | 1 | | GTAGAAGTGGGTTTATTCGTAAAGATGTTA | *boxA* -less: pMME2594 | *boxA* -less: MML918 |
| | 2 | | AGATATCTGCCATATTATACAAACGTTTCC | | |
| | 3 | | CGAGTGACCGGTCTCACACAACTACAATTA | | |
| | 4 | | CACGCCGCATATCGGAATCCACTATCTGCA | | |
| | 5 | | ctctttcgtttattatcaagtacttagtcg | | |
| | 6 | | Atcttaactttaataatatttattctgtcg | | |
| | 7 | | Tagttgaagtttctgggaaatatgcgatgt | | |
| | 8 | *lpg0456 (dotO)* | AATCTATATAAGACTCTGTAGTTTGCTTCA | | |
| | 9 | | GTATAACCGCGTACTTGTAGCGTGAGTGTT | | |
| | 10 | | ATAACCGGATATTAATTAGGTTTAAACTAC | | |
| MC-V-dotO | 1 | | GTATAACCGCGTACTTGTAGCGTGAGTGTT | *boxA* -less: pMME2595 | *boxA* -less: MML919 |
| | 2 | | ATAACCGGATATTAATTAGGTTTAAACTAC | *boxA* (-58): pMME2601) | *boxA* (-58): MML925 |
| | 3 | | GTAGAAGTGGGTTTATTCGTAAAGATGTTA | | |
| | 4 | | ACTGTGAGATTCATTTATATTCGAATCACC | | |
| | 5 | | AGATATCTGCCATATTATACAAACGTTTCC | | |
| | 6 | | CACGCCGCATATCGGAATCCACTATCTGCA | | |
| | 7 | | ctctttcgtttattatcaagtacttagtcg | | |
| | 8 | | Atcttaactttaataatatttattctgtcg | | |
| | 9 | | Tagttgaagtttctgggaaatatgcgatgt | | |
| | 10 | *lpg0456 (dotO)* | AATCTATATAAGACTCTGTAGTTTGCTTCA | | |
| MC-VI-dotO | 1 | | Tagttgaagtttctgggaaatatgcgatgt | *boxA* -less: pMME2596 | *boxA* -less: MML920 |
| | 2 | *lpg0456 (dotO)* | AATCTATATAAGACTCTGTAGTTTGCTTCA | | |

| | | | | | |
|---|---|---|---|---|---|
| | 3 | | GTATAACCGCGTACTTGTAGCGTGAGTGTT | | |
| | 4 | | ATAACCGGATATTAATTAGGTTTAAACTAC | | |
| | 5 | | GTAGAAGTGGGTTTATTCGTAAAGATGTTA | | |
| | 6 | | ACTGTGAGATTCATTTATATTCGAATCACC | | |
| | 7 | | CGAGTGACCGGTCTCACACAACTACAATTA | | |
| | 8 | | CACGCCGCATATCGGAATCCACTATCTGCA | | |
| | 9 | | ctctttcgtttattatcaagtacttagtcg | | |
| | 10 | | Atcttaactttaataatatttattctgtcg | | |
| MC-VII-dotO | 1 | | Tagttgaagtttctgggaaatatgcgatgt | *boxA* -less: pMME2597 | *boxA* -less: MML921 |
| | 2 | *scrambled dotO* | TAATAGGCTCTTCGAATTTATACTCAGTAT | | |
| | 3 | | GTATAACCGCGTACTTGTAGCGTGAGTGTT | | |
| | 4 | | ATAACCGGATATTAATTAGGTTTAAACTAC | | |
| | 5 | | GTAGAAGTGGGTTTATTCGTAAAGATGTTA | | |
| | 6 | | ACTGTGAGATTCATTTATATTCGAATCACC | | |
| | 7 | | CGAGTGACCGGTCTCACACAACTACAATTA | | |
| | 8 | | CACGCCGCATATCGGAATCCACTATCTGCA | | |
| | 9 | | ctctttcgtttattatcaagtacttagtcg | | |
| | 10 | | Atcttaactttaataatatttattctgtcg | | |
