## Supplemental Table 3 for "A multiplex CRISPR interference tool for virulence gene interrogation in an intracellular pathogen"

### Table S3: Primers used in this study

| Cloning Primer | Sequence 5' to 3' |
|---|---|
| BKMP108 | GGGG ACA ACT TTT CTA TAC AAA GTT GTA GACGTCTTAAGACCCACTTTC |
| BKMP109 | GGGG AC AAC TTT ATT ATA CAA AGT TGT CCGACAAACAACAGATAAAAC |
| BKMP94 | GGGG ACA AGT TTG TAC AAA AAA GCA GGC TTA CTTTAGGGCCATTAAAAGTTC |
| BKMP95 | GGGG AC AAC TTT GTA TAG AAA AGT TGG GTG ATGCTCCAACAATTGCAAATATG |
| BKMP98 | GGGG ACA ACT TTG TAT AAT AAA GTT GTA CCCGCAATTAAAGCGCCT |
| BKMP99 | GGGG AC CAC TTT GTA CAA GAA AGC TGG GTA CAGGAAGCCCGGATTGATTG |
| BKMP185 | GGGGACAACTTTGTATACAAAAGTTGTA tatttcttaataactaaaaatatggtataa |
| BKMP186 | GGGGACCACTTTGTACAAGAAAGCTGGGTT aactcaacaagtctcagtgtgctgaa |
| BKMP45 | GGGGACAAGTTTGTACAAAAAAGCAGGCTTA CCATAATGAGTTTGATGATTTCAATAATAG |
| BKMP46 | GGGGACAACTTTTGTATACAAAGTTGT CCTGTGGAGCTTAGTAGGTTTAGCAAGATGGC |
| crsidF_F | taggcg GGTCTC t AAAC GTTCCAGATCACTTAATAAATGCGTGACAT |
| crsidD_F | taggcg GGTCTC t AAAC AGATTTCACTATTACCCTTATCTAAATCAT |
| crsetA_F | taggcg GGTCTC t AAAC TTTGTCAATTTGAGCCTCTTGACCAGCCTG |
| crlidA2_R | gggaca GGTCTC a AAAAc ATTAATGGCCGCATTGTTTTTGTGGTATTT |
| Gateway5-Ptet | GGGG ACA ACT TTG TAT ACA AAA GTT GTA tccctatcagtgatagagattgac |
| Gateway2-T1term | GGGG AC CAC TTT GTA CAA GAA AGC TGG GTTCAG GAG AGC GTT CAC C |
| Gateway5-leader | GGGG ACA ACT TTG TAT ACA AAA GTT GTA AAT AAA TGC AGT AAT ACA GGG GC |
| Gateway2-leader | GGGG AC CAC TTT GTA CAA GAA AGC TGG GTT CTC GTA GAC TAT TTT TGT CTA AAA AAT TTC |
| BoxA_srtF | caagactgaaAtTtagTtCTTTAaaaATTtgcgattacg |
| BoxA_srtR | cgtaatcgcaAATtttTAAAGaActaAaTttcagtcttg |
| BoxA_farF | gagcacAataaatTTagtTCtTTaAAAATttttcaagactg |
| BoxA_farR | cagtcttgaaaaATTTTtAAaGAactAAatttatTgtgctc |

| qPCR Primer | Sequence 5' to 3' |
|---|---|
| ctrl_rpsL_F | GCTATGCGTAAGGTTGCACG |
| ctrl_rpsL_R | GCGCACACCAGGTAAATCCT |
| RT_2815_F | AGCTACTTGACTGGCTTGCT |
| RT_2815_R | TTCCCAGCCCCATGAAAAGG |
| RT_lidA_F | ACCCTTCTCGGCCCAGATTA |
| RT_lidA_R | CAATGAGCCAGTCAAGCCCT |
| RT_vipD_F | CCCATGTTAGCGGAGCATCT |
| RT_vipD_R | CTCCTCTCGCACGAAAACCT |
| RT_sidM_F | TGCACCTAAAGAGGGTGCAG |

| | |
|---|---|
| RT_sidM_R | TCCTGGTGCCGAAAGATGAG |
| RT_dotO_F | TGATTGGCGCGCAACTTTAC |
| RT_dotO_R | CAAAACCTGCCTGGACAACG |
| RT_dotD_F | CTTGCAAGCAAGAGCCAGTG |
| RT_dotD_R | GGCGACTTGCCTAAAACACG |
| RT_ravN_F | TGGGCACGAGAGGTACTACA |
| RT_ravN_R | TCCTGTTGTTGAGCAGGGTT |
| RT_lpg2793_F | CTCTTCGCCGGCCGATTT |
| RT_lpg2793_R | CCCAGAGCAGGCATTTCTTG |
| RT_lpg0208_F | TGGATGCGGATATGCAGTGT |
| RT_lpg0208_R | CGCTAGCCTGAGAGAAAGCA |
| RT_lpg1356_F | ACAAGGCTATAGCAAGGCACA |
| RT_lpg1356_R | TTTTCATCCCCCTGCTCTGC |
| RT_lpg0140_F | TATCGTTGACGCAGGCCATT |
| RT_lpg0140_R | GCAGCACGACAAAATCACGA |
| RT_lpg0246_F | ATGCTCTGTATGGGTGAGGC |
| RT_lpg0246_R | ACCAGGCATTTAACCGTTGC |
| RT_lpg2733_F | GCAGCGGGATGCATTGTTTT |
| RT_lpg2733_R | CAACCATCCCCATCCTGTGG |
